# Particle-associated and free-living microbial assemblages are distinct in a permanently redox-stratified freshwater lake

**DOI:** 10.1101/2021.11.24.469905

**Authors:** Ashley B. Cohen, Vanja Klepac-Ceraj, Kristen Butler, Felix Weber, Arkadiy I. Garber, Lisa N. Christensen, Jacob A. Cram, Michael L. McCormick, Gordon T. Taylor

## Abstract

Microbial assemblages associated with biogenic particles are phylogenetically distinct from free-living counterparts, yet biogeochemically coupled. Compositions may vary with organic carbon and inorganic substrate availability and with redox conditions, which determine reductant and oxidant availability. To explore microbial assemblage compositional responses to steep oxygen and redox gradients and seasonal variability in particle and substrate availability, we analyzed taxonomic compositions of particle-associated (PA) and free-living (FL) bacteria and archaea in permanently redox-stratified Fayetteville Green Lake. PA and FL assemblages (> 2.7 *µ*m and 0.2 – 2.7 *µ*m) were surveyed at the peak (July) and end (October) of concurrent cyanobacteria, purple and green sulfur bacteria blooms that result in substantial vertical fluxes of particulate organic carbon. Assemblage compositions varied significantly among redox conditions and size fractions (PA or FL). Temporal differences were only apparent among samples from the mixolimnion and oxycline, coinciding with seasonal hydrographic changes. PA assemblages of the mixolimnion and oxycline shifted from aerobic heterotrophs in July to fermenters, iron-reducers, and denitrifiers in October, likely reflecting seasonal variability in photoautotroph biomass and inorganic nitrogen. Within a light-scattering layer spanning the lower oxycline and upper monimolimnion, photoautotrophs were more abundant in July than in October, when *Desulfocapsa*, a sulfate-reducing and sulfur-disproportionating bacterium, and Chlorophyte chloroplasts were abundant in PA assemblages. In this layer, microbial activity and cell concentrations were also highest. Below, the most abundant resident taxa were sulfate-reducing bacteria and anaerobic respirers. Results suggest PA and FL assemblage niche partitioning interconnects multiple elemental cycles that involve particulate and dissolved phases.

## Introduction

Biogenic debris can be rich in organic matter, inorganic nutrients, and minerals and teem with microbes that extract those resources more efficiently than free-living microbes in surrounding, nutrient-depleted waters. (Smith et al., 1992, Taylor et al., 2009). As microbes carry out remineralization or photosynthesis in particles, they generate byproducts such as dissolved organic matter and inorganic nutrients that diffuse out to free-living microbes in the vicinity of the particles over spatial scales up to 100 times larger than the particle itself (Kiørboe et al., 2001). This interplay between the particle-associated (PA) and free-living (FL) microbial assemblages intensifies seasonally during algal blooms, when chemical gradients emanating from algal-rich particles (the phycosphere) support FL microbes that consume algal exudates (Cai et al., 2014, Louati et al., 2015; Kieft et al., 2021).

In permanently redox-stratified meromictic lakes, marine basins, marine oxygen-deficient zones, and many fjords, settling particles pass through distinctive redox zones with correspondingly distinctive FL microbial assemblages as they transit towards the underlying sediment by gravitational settling (Fuchsman et al., 2011, Suter et al., 2018). Therefore, the interplay between microbes in particles with resident free-living microbes likely differs with redox conditions due to the changing oxidant and reductant availability. In these quiescent, diffusion-dominated environments, redox conditions can vary with both depth and particle attributes. Owing to minimal advective chemical exchange between particles and water (Alldredge and Cohen, 1987, Plough et al., 1997, Kiørboe et al., 2001), aerobic respiration, fermentation, denitrification, sulfate reduction, methanogenesis, and anaerobic methane oxidation can all be supported simultaneously in oxyclines existing at spatial scales of micrometers to meters (Fuchsman et al., 2011, Lauro et al., 2011, Wright et al., 2012, Ganesh et al., 2015, Suter et al., 2018, Torres-Beltrán et al., 2019).

To explore how ambient redox conditions and seasonally varying organic carbon substrates influence the interplay between PA and FL microbes, we used permanently redox-stratified (meromictic), sulfidic, Fayetteville Green Lake (FGL), NY, USA (43°03′32.3″N, 75°58′18.4″W) as a natural laboratory. FGL is one of the most extensively studied meromictic lakes, and has served as a natural laboratory to study many aspects of biogeochemistry in permanently redox-stratified water columns (e.g., Zerkle et al., 2010, Meyer et al., 2011, Havig et al., 2015, 2018, Fulton et al., 2018). FGL has unusually high sulfate concentrations (∼15 mM) for a freshwater environment because groundwater entering near the upper oxycline boundary, and potentially in the deep lake, passes through gypsum-bearing shale (CaSO_4_ · 2H_2_O) (Brunskill and Ludlam, 1969, Havig et al., 2015). The high sulfate concentrations and permanent density stratification result in vertical biogeochemical zonation similar to marine anoxic basins, such as the Cariaco Basin and Black Sea. Zones include a fully oxygenated mixed layer (mixolimnion) and a sulfidic deep layer (monimolimnion) separated by an oxycline within which dissolved O_2_ concentrations attenuate steeply until hydrogen sulfide first appears. FGL’s mixolimnion (0-15 m) is normoxic (O_2_ ≥ 93 *µ*M), while the oxycline (15-20 m) varies from oxic to hypoxic (93 > O_2_ ≥ 3 *µ*M) and suboxic (O_2_ < 3 *µ*M, no measurable H_2_S), and the monimolimnion (≥ 20 m) is euxinic (measurable H_2_S) Havig et al. (2015, 2018) as defined in Scranton et al. (2014) and Taylor et al. (2018).

Photoautotrophic populations in FGL have been extensively studied for more than 50 years. These include cyanobacteria in the mixolimnion and near the lower oxycline boundary that induce calcium carbonate precipitation (“whiting events”), and populations of purple and green sulfur-oxidizing anoxygenic photoautotrophs at and just below the lower oxycline boundary (Culver and Brunskill, 1969, Thompson et al., 1990, 1997, Fulton et al., 2018, Block et al., 2021). They bloom during July, when particulate organic carbon fluxes to the deep lake are elevated, and senesce during October, when particulate organic carbon fluxes to the deep lake decrease (Brunskill, 1969, Culver and Brunskill, 1969), suggesting that they are a significant source of organic carbon to anoxic waters and to the lakebed.

Less is known about the non-photoautotrophic members of FGL microbial assemblages and their roles in biogeochemical cycling. Interactions between photoautotrophs and putative chemoautotrophic sulfur-disproportionating bacteria in the oxycline and upper monimolimnion (15-21 m) and between chemoorganotrophic sulfate-reducing and fermentative bacteria in the shallow monimolimnion (22-25 m) couple the lake’s complex carbon and sulfur cycles (Block et al., 2021). In the deeper monimolimnion (21-52 m), organic substrates (e.g., chitin, lignin, cellulose, acetate, butyrate) appeared to be more influential on assemblage composition than electron acceptors in enrichment incubations, suggesting that the niche-partitioning of organotrophs drives its structure and function (Rojas et al., 2021). Complex, higher molecular weight organic substrates are likely delivered to the deeper monimolimnion by gravitational settling of biogenic particles from the water column above, while monomeric organic substrates are likely produced in situ from anaerobic diagenetic processes. Photoautotrophic blooms appear to be the major source of POC-associated substrates, as 74% of the lake’s annual POC flux to the deep monimolimnion occurs during the bloom period (calculated from Brunskill, 1969).

Biogenic particles seem to be major foci of coupled biogeochemical cycles and interactions between primary producers and chemoorganotrophs in FGL. However, to our knowledge, the identities of microorganisms populating PA and FL assemblages in this system are still not known. At the least, their identities are needed to infer the functions of and synergies between PA and FL assemblages and how they affect biogeochemical cycling in FGL. These inferred functionalities can be developed into testable hypotheses and may aid in building models of biogeochemical cycling. Furthermore, few studies characterizing PA and FL assemblages in shallow, permanently redox-stratified water bodies with intensive blooms of anaerobic photoautotrophs (e.g., meromictic lakes, fjords, Lauro et al., 2011, Torres-Beltrán et al., 2019) exist. To date, such studies have been conducted primarily in large-scale marine anoxic basins (Fuchsman et al., 2011, Suter et al., 2018, Suomenin et al., 2020), despite the key role played by lakes and oxygen-depleted waters in global carbon sequestration and climate change (Downing et al., 2008, Brietburg et al., 2015).

Here, we examine prokaryotic PA and FL assemblages in FGL throughout all redox zones. Our goal is to better understand the roles of PA microbes in biogeochemical processes under varying redox conditions in freshwater systems. We characterize how PA and FL assemblages organize throughout the FGL water column using 16S rRNA gene amplicon sequences and geochemical/hydrographic profiling. We sampled during the peak of concurrent cyanobacteria and purple and green sulfur bacteria blooms in July and at their termination in October to gain some insight into how seasonal surface processes affect community structure and biogeochemical cycles. We hypothesize that redox conditions, bloom events, and particle formation (assemblage type) are the strongest determinants of community structure and activity. Through this work, we hope to highlight the importance of biogenic particles in biogeochemical cycling in FGL and other permanently redox-stratified freshwater environments and build hypotheses for more targeted future studies.

## Materials and methods

### Sample and sensor data collection

We conducted field work on 5-7 October 2016 and 17-21 July 2017 above the deepest part of the lake (43°03′01.9″N, 75°57′58.9″W, ∼52 m deep) from a rowboat and a pontoon boat. Sampling depths spanned the entire water column and strategically targeted the lower oxycline through upper euxinic zone, where physico-chemical features are narrowly distributed (Havig et al., 2015, 2018), at finer (meter to sub-meter) vertical resolution (Table 1). To determine the basic physico-chemical structure and redox-zonation of the water column, a suite of continuous sensor data measurements and hydrogen sulfide concentration measurements were made. Concentrations of oxygen, phycoerythrin and chlorophyll-a, as well as turbidity, salinity, oxidation-reduction potential (ORP), total dissolved solids, and pH were all continuously profiled using a YSI EXO1 sonde sensor package. Discrete samples for hydrogen sulfide concentration measurements and DNA recovery and analyses were collected using Tygon tubing held straight with a planar weight attached to the sampling end and connected to a peristaltic pump. Hydrogen sulfide samples (10 mL) were collected in triplicate by immersing a glass gas-tight syringe in an open 60 cc syringe body continuously flushed with water from the collection depth. The glass syringe was purged several times, and contents were immediately expelled into vial with 1 mL of 1M zinc acetate preserving dissolved sulfide as ZnS. Samples were refrigerated in the dark until analysis (Li et al. 2008, Zerkle et al., 2010, Li and Astor, 2011). Samples for prokaryotic cell counts (50 mL) were collected in duplicate at each depth, immediately fixed with freshly 0.2 *µ*m filtered borate-buffered formaldehyde (2% final conc.), and frozen on dry ice. Samples were stored at −20°C upon returning to Stony Brook University, NY, USA.

**Table 1.** Discrete sample analyses performed per sampling depth during July and October. N/A=not applicable, depth not sampled. “All” indicates that samples were recovered for 16S rRNA gene amplicon sequencing, hydrogen sulfide concentration measurements, total prokaryote cell counts, bacterial heterotrophic production rate measurements, and inorganic carbon assimilation rate measurements.

|  | 5-7 Oct. 2016 | 17-21 Jul. 2017 |
| --- | --- | --- |
| Depth (m) |  |  |
| 5 | all | N/A |
| 10 | N/A | all |
| 15 | all | all |
| 16 | only hydrogen sulfide and 16S rRNA amplicons | all |
| 19 | only hydrogen sulfide and 16S rRNA amplicons | all |
| 20 | all | all |
| 20.5 | N/A | all |
| 21 | only hydrogen sulfide and 16S rRNA amplicons | all |
| 23 | N/A | all |
| 25 | all | all |
| 40 | all | all |

Samples reserved for DNA recovery and analyses were separated into particle-associated (PA) and free-living (FL) operationally-defined fractions by filtering the one liter of lakewater collected from each sampling depth sequentially through sterile 2.7 *µ*m QMA and 0.2 μm Sterivex™ in-line filters using a peristaltic pump. We recognize that some sequences recovered from the PA fraction may actually be symbiotically-associated with protists or occur in cell aggregates. By the same token, some sequences in the FL fraction may have detached from particles during sample processing. To ensure representation of in situ assemblages, samples were immediately placed on ice in the dark when collected and filtered on shore within 2-3 hours of sampling. Filters were immersed in lysis buffer (40 mM EDTA, pH 8.3 50 mM Tris, 0.73 M sucrose) and immediately snap frozen on dry ice (Suter et al., 2018). Upon returning to the laboratory, samples were stored at −80°C until processing.

### Radiotracer incubations and processing

To measure total autotrophic and bacterial heterotrophic production rates, radiotracer incubation samples were pumped directly into Pierce™ screw-top glass septum vials from each sampling depth. Oxygen contamination was avoided by slowly filling vials from the bottom up, overfilling the vial volume three times and constantly dripping fresh ambient water from the collection depth over the Teflon-lined butyl rubber septum while sealing (Taylor et al., 2001). Vials were placed on ice in the dark until incubations were started in 1-3 hours after sampling.

Inorganic carbon assimilation radiotracer samples were amended with N_2_-purged ^14^C-labelled bicarbonate (Moravek Biochemicals) to a final activity of 0.045 *µ*Ci/mL, which equates to an addition of 7.76 x 10^-4^ mM radioactive bicarbonate to 5-10 mM ambient FGL dissolved inorganic carbon pools (Havig et al., 2018). Vials were incubated for 24 h and terminated with orthophosphoric acid (0.03 N final concentration) (Taylor et al., 2001).

To measure the total bacterial heterotrophic production, samples were amended with N_2_-purged ^3^H-leucine (MP Biomedicals), a representative labile dissolved organic carbon substrate, to a final activity of 0.016 *µ*Ci/mL. The working solution was created by adding radioactive leucine to a cold leucine carrier such that the final leucine concentration in incubations was 22 nM. This leucine concentration is believed to be sufficient to saturate bacterial uptake systems and therefore measure potential assimilation rates in FGL, which is oligotrophic (approximately half the concentration required for uptake saturation in eutrophic lakes, Bastviken and Tranvik, 2001). Vials were incubated for 8 h and terminated with trichloroacetic acid (5% TCA w/w final concentration) (Kirchman 1993, Taylor et al. 2001).

Sample vials and TCA-killed controls were incubated in mesh bags layered to transmit 5.4, 3.2 and 2.5% photosynthetic active radiation (PAR) to mimic in situ PAR and were placed in open water incubators (one for each FGL temperature zone) to maintain in situ temperatures (**Supplementary materials**, Fig. S4). Bags were immersed on floating racks to maintain desired temperatures and illumination. Vials from aphotic depths were placed in opaque dark bags at the bottom of the incubator. Temperatures were maintained by adding ice or warm water as needed. In situ PAR was calculated as in Taylor et al. (2003) using published July Secchi depths (Culver and Brunskill, 1969).

Terminated inorganic carbon assimilation samples were vacuum filtered (≤ 200 mm Hg) onto 0.2 *µ*m nitrocellulose filters (25mm diam. GVS S.p.A) to dryness. Filters were then rinsed twice with chilled, freshly filter-sterilized, acidified lake water (adjusted to pH 3.5 with H_3_PO_4_) to remove unincorporated ^14^C-bicarbonate. To further remove unincorporated ^14^C-bicarbonate, filters were fumigated in their glass scintillation vials within a sealed basin containing an open vial of concentrated (12N) HCl for 24 h and then air-dried.

Terminated bacterial heterotrophic production samples were maintained at 80°C for 20 min in vented test tubes to precipitate proteins, then chilled before further handling. Precipitated proteins were captured on 0.2 *µ*m nitrocellulose filters (25mm diameter) by vacuum filtration (≤ 200 mm Hg). Filters were rinsed twice with 3 ml of chilled 5% TCA, twice with 3 ml of chilled 80% ethanol, then saturated dropwise with a final 1 ml 80% ethanol rinse. Filters were dissolved with 0.5 ml ethyl acetate in glass scintillation vials (Kirchman, 1993).

During each round of bacterial heterotrophic production and inorganic carbon assimilation filtrations, killed controls for abiotic sorption or precipitation of labelled materials were filtered after all environmental samples. For specific activity calculations, the same tracer volume inoculated into samples was added to a small volume of filter-sterilized lake water. These solutions account for how much radioactive tracer is available for biological uptake once accounting for matrix effects. Sample membranes and specific activity solutions were then radioassayed by liquid scintillation counting. ^14^C-bicarbonate radioactivity counts (disintegrations per minute, dpm) were converted to inorganic carbon assimilation rates as in Taylor et al. (2001) using published dissolved inorganic carbon concentrations for FGL from Havig et al. (2015). ^3^H-leucine radioactivity dpm were converted to bacterial heterotrophic production according to Kirchman (1993) and Taylor et al. (2001).

### Hydrogen sulfide sample processing

We used Cline’s method (Cline, 1969) to quantify hydrogen sulfide. Hydrogen sulfide standards were made by adding Na_2_S*9H_2_O to de-aerated (boiled) nitrogen-purged ddH_2_O and diluted to produce 7-8-point calibration curves. Standards were preserved in the same manner as field samples. Prior to analysis, samples, blanks, and standards were allowed to come to room temperature in the dark and split equally into two Falcon tubes containing 44.5 mL of ddH_2_O each. In this study, 2 ml of diamine reagent was added to each Falcon tube, sample color development proceeded for 30 min in the dark at room temperature, and absorbance at 665 nm was measured spectrophotometrically.

### Prokaryotic cell enumeration

To obtain total prokaryotic cell counts, subsamples (2 - 5 ml) were stained with 4′,6-diamidino-2-phenylindole (DAPI), then filtered through 0.2 *µ*m black polycarbonate membranes (25 mm diam. Whatman Nuclepore™) according to standard protocols (Porter and Feig 1980). To optimize cell dispersion, membranes were dipped in freshly filtered (0.2 *µ*m) 0.01% Triton X-100 (Sigma-Aldrich) detergent and mounted on water-saturated nitrocellulose backing filters on metal vacuum filtration frits. Membranes were air dried and mounted with a 4:1 mixture of Citifluor™ (Ted Pella, Inc.) and Vectashield® (Vector Laboratories Inc.) mounting solutions. Cells were enumerated on a Zeiss Axioscope epifluorescence microscope at 1000x magnification among sufficient fields of view to achieve ≤ 10% relative standard deviation (typically between 10 and 20 fields of view).

### DNA extraction and amplicon sequencing

To recover high-quality genomic DNA for Illumina 16S rRNA amplicon sequencing, DNA was extracted, purified, and concentrated as in Suter et al. (2018). Briefly, sample and negative control lysates were sequentially extracted with agitation using Lysozyme (Fisher Bioreagents) and SDS combined with Proteinase K (Fisher Bioreagents, Sigma-Aldrich®) enzyme at elevated temperature (37 °C and 55°C, respectively), followed by pH 8-buffered 25:24:1 phenol:chloroform:isomyl alcohol and 24:1 chloroform:isomyl alcohol (Sigma-Aldrich®). Extracts were concentrated using 100 kDa membrane centrifugation (Amicon®), and then further concentrated and purified using a Zymo Genomic DNA Clean & Concentrator® kit. Quality-indicating A_260/280_ ratios and double-stranded DNA concentrations of purified DNA samples were measured in triplicate using a NanoDrop Lite (Thermo Scientific).

To further assess DNA quality and possible inhibition prior to sequencing, polymerase chain reactions (PCRs) were prepared using the primers Bac331F (5’-TCCTACGGGAGGCAGCAGT-3’) and Bac797R (5’-GGACTACCAGGGTATCTAATCCTGTT-3’), which amplify the bacterial 16S rRNA gene (Nadkarni et al. 2002). For each undiluted sample, 10-fold and 100-fold dilutions prepared with nuclease-free water were used as templates in separate PCR reactions prepared with Lucigen Failsafe™ reagents. Reactions using Pre-mix J and 600 nM forward and reverse primer were prepared according to the manufacturer’s instructions. All reactions were run along with a no template control (NTC) and a full procedural blank using a LabNet MultiGene™ Optimax thermal cycler. Reactions were subjected to a pre-denaturation stage of 5 min at 95°C for 1 cycle, a 40 cycle PCR stage consisting of 0.5 min denaturation at 95°C, annealing at 50°C for 0.5 min, and extension at 72°C for 1.5 min per cycle, and a final 1 cycle extension stage at 72°C for 15 min. PCR products were qualitatively evaluated using a UVP GelDoc-It® system after gel electrophoresis using a 1% agarose gel.

Bacterial and archaeal community composition, diversity, and assemblage-partitioning was assessed by sequencing amplicons of the V4-V5 hypervariable regions of the 16S rRNA gene. We used the universal primers 515FY (5’-GTGYCAGCMGCCGCGGTAA-3’) and 926R (5’-CCGYCAATTYMTTTRAGTTT-3’) (Parada et al., 2016). These primers had suitable Illumina adapters and unique barcodes, allowing all samples to be run simultaneously in a single flow cell. All samples were prepared for sequencing following the Microbiome Amplicon Sequencing Workflow (Comeau et al., 2017). Amplicons were amplified in duplicate samples from the 1:1 and 1:10 DNA template dilution with the exception of 20 and 20.5 m July samples, which were amplified from the 1:10 and 1:100 DNA template dilutions due to inhibitively high DNA concentrations. Paired-end read sequencing (∼350 bp) was performed by the Integrated Microbiome Resource (Dalhousie U., Halifax, Nova Scotia, Canada) using an Illumina MiSeq sequencing platform, producing on average 44,000 raw reads per sample. Raw sequence files were deposited to the NCBI Sequence Read Archive under the BioProject accession number PRJNA752637.

### Processing sequence data

Taxonomic assignments of amplicon sequence variants (ASVs), ASV counts per sample, the relative abundances of taxa at specified taxonomic levels, analysis of microbial community (ANCOM) statistics, and alpha (mean species) and beta diversity (community differentiation) matrices were generated using QIIME 2™ version 2020.2 (Boylen et al., 2019), based on the LangilleLab workflow (Comeau et al., 2017) and QIIME 2™ moving pictures tutorials with slight modifications, as described below.

First, the raw sequences (demultiplexed Casava 1.8 format) were prescreened for adapter sequence contamination, min and max read length and low-quality reads and these were removed prior to trimming primers from the reads. Read files were then further examined using FastQC (Andrews, 2010) to determine the portion of forward and reverse reads retained during read truncation.

Using the QIIME 2™ plugin ‘q2-cutadapt trim-paired,’ primer sequences were trimmed from reads and all reads that did not begin and end with the forward and reverse primer sequences were removed (Martin, 2011). Trimmed reads were truncated and denoised with DADA2 using the ‘q2-dada2’ plugin (Callahan et al., 2016) with the allowable base-call error for forward and reverse reads set to 3, which retained an average of 23,078 non-chimeric reads per sample. The resulting table of sequence (QIIME 2™ “feature”) frequency per sample from the DADA2 output was then frequency-filtered to retain sequences with a minimum frequency of 20 reads in at least one sample to exclude features that likely occurred due to MiSeq bleed-through between runs.

The frequency-filtered sequence frequency table was then used to filter the sequence table from the DADA2 output to create a table of representative ASVs. Those ASVs were classified using ‘q2-classifier’ plugin (Bokulich et al. 2018) with a classifier that was pre-trained against the SILVA database for our primers (silva_132_99_16S_V4.V5_515F_926R database) using the Naive-Bayes approach implemented in the ‘scikit-learn’ Python library. The classified frequency-filtered frequency table was then filtered by taxonomy to exclude chloroplasts and mitochondria using the “taxa filter-table” command. The removed chloroplast and mitochondria sequences that did not belong to cyanobacteria were run through megaBLASTN^+^ to identify eukaryote algae according to the accepted definition of an operational taxonomic unit (>97% amplicon percent identity) (Altschul et al., 1990, Morgulis et al., 2008, Camacho et al., 2008). Steps preceding filtration by taxonomy were then repeated with the taxonomy-and-frequency filtered sequence frequency table to produce the final sequence and taxonomy tables. Counts per sample tables were output in both the ASV format and with ASVs collapsed into taxa at desired taxonomic levels by exporting the taxonomy-filtered feature frequency table or using the “taxa collapse” command.

The mitochondria-and-chloroplast free ASVs were then aligned with the multiple alignment program for amino acid or nucleotide sequences (MAFFT, Katoh et al., 2002) using default settings. Aligned ASVs were used to construct a phylogenetic tree with fasttree2 (Price et al, 2010) using default settings through the ‘q2-phylogeny’ plugin. To ensure that sequencing depth fully saturated the alpha diversity of samples with a lower number of reads, a rarefication plot of various alpha diversity metrics versus reads was produced by rarifying to 4900 sequences (the minimum number of reads out of all samples) per sample with a step size of 20 sequences. Using the ‘q2-diversity’ plugin, we calculated alpha diversity indices, Shannon’s Diversity Index (H) that accounts for both taxon abundance and evenness (Shannon and Weaver, 1949) and Pielou-evenness that assesses how evenly taxa are distributed within the community (Pielou, 1966), and beta diversity indices, Bray-Curtis that quantifies compositional similarities between two communities (Bray and Curtis, 1957) and weighted unifrac that quantifies compositional similarities and incorporates phylogenetic distances of taxa within the communities (Lozupone et al., 2007).

To determine if any ASVs were significantly more abundant in assemblages defined by metadata categories, <u>an</u>alysis of microbial <u>com</u>munity (ANCOM) (Mandal et al., 2015) was performed. A pseudo-counted chloroplast-and-mitochondria-free filtered ASV table was subject to ANCOM using the QIIME 2™ “composition ancom” command; the pseudo-count increases all ASV counts by one because zero is not allowed as input. ANCOM tests determine if ASVs’ relative abundances in two assemblages are significantly different by calculating pairwise log relative abundance ratios between all representative ASVs in each assemblage. Per ASV, from these pairwise ratios, a W-statistic (the number of ratios that are significantly different between the two assemblages by an f one-way test) is calculated. The relative abundance of the ASV in the two assemblages assemblages is considered significantly different if the W-statistic is above an empirical threshold. To ensure that significant W-statistics were not due to the ASV having a higher relative abundance than most ASVs in a given sample, ANCOM test results were examined in clr (the log_2_-transformed arithmetic mean-normalized relative abundance)-W statistic space. Then only significant ASVs that were not over-represented (in the upper left-hand quadrant of the volcano plot) were retained.

All correlation analyses between alpha and beta diversity metrics and metadata were performed using a categorical version of the metadata. A categorical metadata table was created by binning all continuous metadata measurements after histogram visualization other than redox zones, which were defined as presented in the introduction.

The relative <u>o</u>verrepresentation <u>f</u>actor (OF) of a taxon in the particle-associated fraction of a given sample was calculated by:

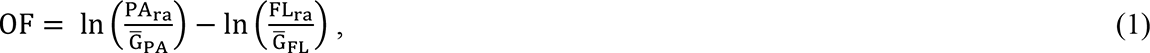

where 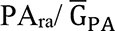 is the relative abundance of that taxon in the particle-associated assemblage normalized to that particle-associated sample’s relative abundance geometric mean. 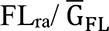 is the relative abundance of the same taxon in the free-living assemblage normalized to that free-living sample’s relative abundance geometric mean. A taxon was considered over-represented in the particle-associated or free-living assemblage if their OF was 1.25 or −1.25, respectively. These bounds correspond to a 3.5-fold difference.

This metric, based on log-transformed geometric mean-normalized relative abundances, was created in lieu of a standard enrichment factor (log-transformed 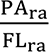) to allow true between-sample numeric comparisons. Log-transformed geometric mean-normalized relative abundances, also known as “center log ratio (clr) transformed,” are commonly used in microbiome analyses. Clr-transformation allows for scale-invariant between-sample comparisons regardless of the abundance distributions of taxa in individual samples, making between-sample statistics more meaningful (Gloor et al., 2017, Badri et al., 2018). As an example, clr-transformed abundance data is commonly plotted against test statistic data on volcano plots as part of ANCOM analyses to ensure that ASVs are not returned as significant due to skewed abundance data (e.g., Cendron et al., 2020, Li et al., 2020).

All alpha and beta diversity statistics other than the dissimilarity matrices were calculated using the python sci-kit bio™ library (v.0.5.6). Principal coordinate analysis (PCoA) was performed on both Bray Curtis and weighted unifrac beta QIIME 2™ dissimilarity matrices. This produces principal coordinates from combinations of the matrix such that new variables are uncorrelated and most matrix variance is found in the first two or three variables. Sample clustering in PCoA matrices was visualized in the three-dimensional space of the first three principal coordinates. Samples were color-coded by categorical metadata to match categories to clustering patterns. Bray-Curtis and weighted unifrac visualizations were compared to determine if incorporating phylogenetic tree branch lengths improved pattern matching between metadata categories and clustering patterns. Permutational analysis of variance (PERMANOVA) was performed on beta diversity matrices to determine if metadata variables explained dissimilarity among samples. PERMANOVA compares the ratio of the sum of squares of a chosen distance metric (in this case, Bray-Curtis or weighted unifrac dissimilarity) within a group to the sum of squares between groups to multiple data permutations. An ANOSIM test was run on beta diversity matrices grouped by field campaign month, redox regime, and assemblage type (particle-associated or free-living). The null hypothesis of an ANOSIM test is that similarities among samples within a particular category/grouping are less than or equal to similarities among samples in the other categories/groupings. The test statistic R is calculated as the ratio of the difference between the average rank similarity between sample pairs from within categories and between categories. R can be anywhere from −1 to 1, with positive values indicating similarity within a category and negative values indicating dissimilarity among categories.

For each metadata variable, Kruskal-Wallis tests were run for each unique categorical bin combination with Pielou evenness values to determine if the variable accounted for significant differences in sample alpha diversity. False discoveries were corrected using the Benjamini/Hochberg (non-negative) method (Benjamini and Hochberg 1995, 2000).

## Results

### Major biogeochemical features

The depth of the wind-mixed, oxygenated layer in the mixolimnion, and the depth of the lower oxycline boundary, defined by the first appearance of H_2_S and precipitous drop in oxidation reduction potential, was shallower during July (3 m and 20 m, respectiveley) than during October (9 m and 21 m, respectively) (Figs. 1, 2, **Supplementary materials** Fig. S5). During both sampling times, the upper oxycline boundary was located at 15 m, and a pronounced light-scattering layer (measured turbidity) was found between 19 and 23 m. H_2_S concentrations in the deep monimolimnion reached maximum concentrations of 1.62 mM (40 m) during July and 2.55 mM (45 m) during October (Fig. 1). These measurements are consistent with previous studies with the exception of the maximum October H_2_S concentration, which was approximately 14% greater than reported values at similar times of the year (Havig et al., 2015, 2018, Fulton et al., 2018, Rojas et al., 2021, Block et al., 2021).

**Figure 1.**
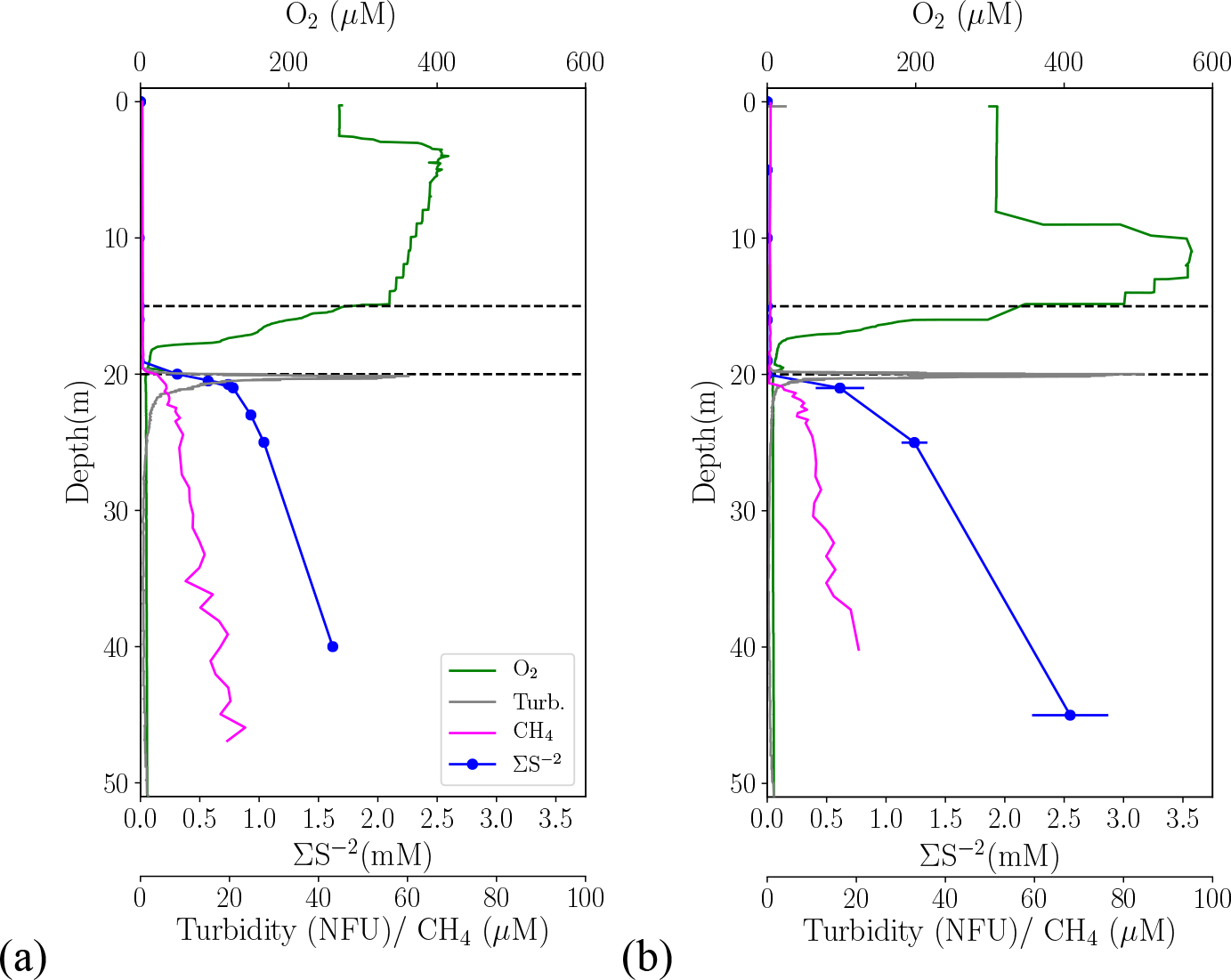
Vertical profiles of major biogeochemical features measured during (a) July 2017 and (b) October 2016, with the exception of methane (CH_4_), which was measured in July 2015 and November 2012 (Havig et al., 2018). Broken lines indicate oxycline boundaries. The standard deviations of sulfide measurements are represented by error bars, but they are too small to be seen.

**Figure 2.**
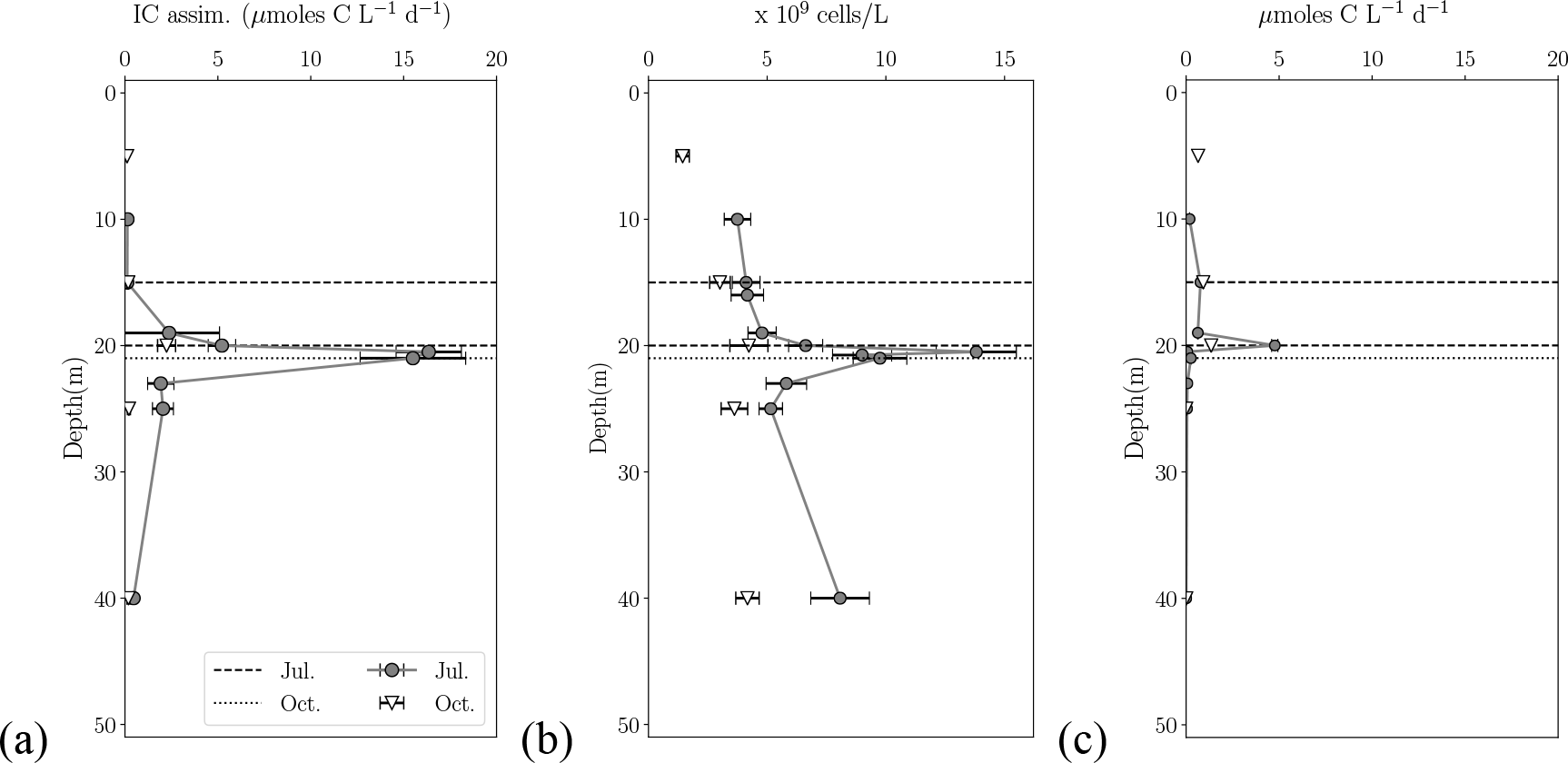
Vertical profiles of (a) inorganic carbon assimilation (ICA) rates, (b) DAPI-stainable cell concentrations (total prokaryotic biomass) and (c) bacterial heterotrophic production (BHP) during July 2017 and October 2016. The lower oxycline boundary is shown as a dotted line for October and as a dashed line for July. The upper oxycline boundary is shown as a dashed line in all plots.

The light-scattering layer had elevated microbial activity and hosted stratified populations of different photoautotrophs during both sampling times. A deep cyanobacterial population was detected by phycoerythrin fluorescence and a deep O_2_ enrichment of approximately 25 *µ*M near the top of the layer (19-19.75 m) in July (Fig. 3). These features overlapped with the maximum rate of bacterial heterotrophic production. Phycoerythrin fluorescence was barely detectable during October, when maximum bacterial heterotrophic production was nearly five-fold lower, deep O_2_ enrichment was 3-fold lower, but chlorophyll-a fluorescence was approximately 3-fold higher (Fig. 3). Brightfield microscopy and observations of intense purple and green water color during both field samplings revealed purple and green sulfur bacteria populations within layers from 20-21 and 21-23 m, respectively. The purple sulfur bacteria population coincided with maxima in transmissometer light attenuation (measured turbidity), total inorganic carbon assimilation rates, total cell concentrations, and pH at or directly below the lower oxycline boundary (Fig. 3**, Supplementary materials** Fig. S6).

**Figure 3.**
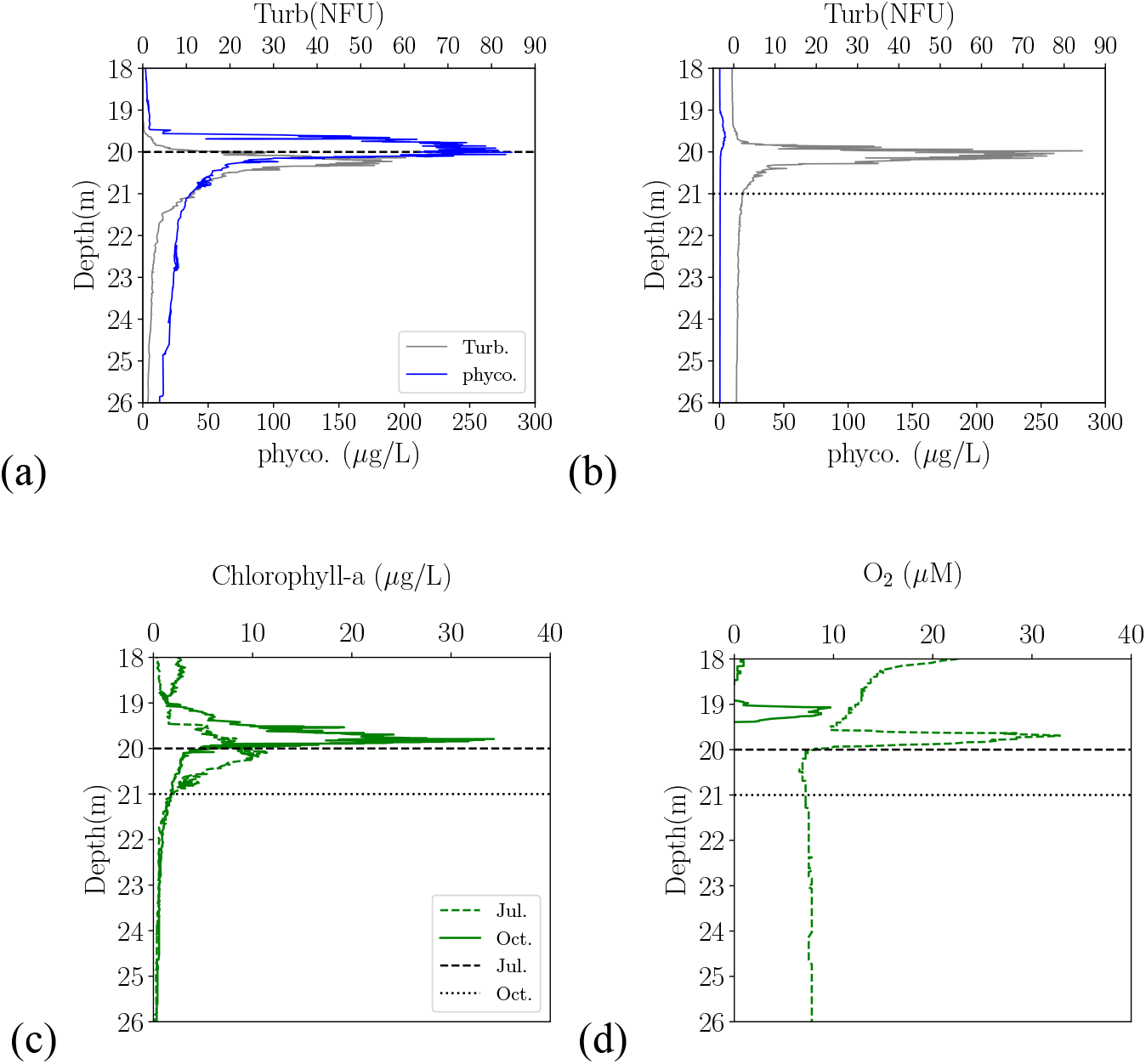
Finer vertical resolution profiles across the lower oxycline and upper monimolimnion of: phycoerythrin fluorescence (cyanobacterial pigment) and turbidity during (a) July 2017 and (b) October 2016. (c) chlorophyll-a fluorescence during July 2017 and October 2016. (D) O_2_ concentration during July 2017 and October 2016. The lower oxycline boundary is shown as a dotted line for October and as a dashed line for July.

### Diversity analyses

Shannon diversity (H) and Pielou evenness alpha diversity indices were used to characterize richness and evenness of the PA and FL assemblages across sampled depths. During October and July, the light-scattering layer (19-23 m) was the least diverse in the water column (Fig. 4). Within this layer, distinctive H minima in PA and FL assemblages were observed during July at 21 m (PA H=4.7, FL H=3.9) and October at 20 m (PA H= 2.6, FL H=5.0), coinciding with purple and green sulfur bacteria and Desulfobulbaceae populations. Pielou evenness index profiles were very similar to the H index, suggesting that low H values are likely caused by a few highly abundant taxa (Fig. 4). When Pielou evenness scores were binned by inorganic carbon assimilation rate, evenness of samples with elevated rates, a characteristic of the light-scattering layer, was significantly different from all other groups (false discovery-corrected pairwise Kruskall-Wallace test *p* < *q*, **Supplementary materials** Table S1).

**Figure 4.**
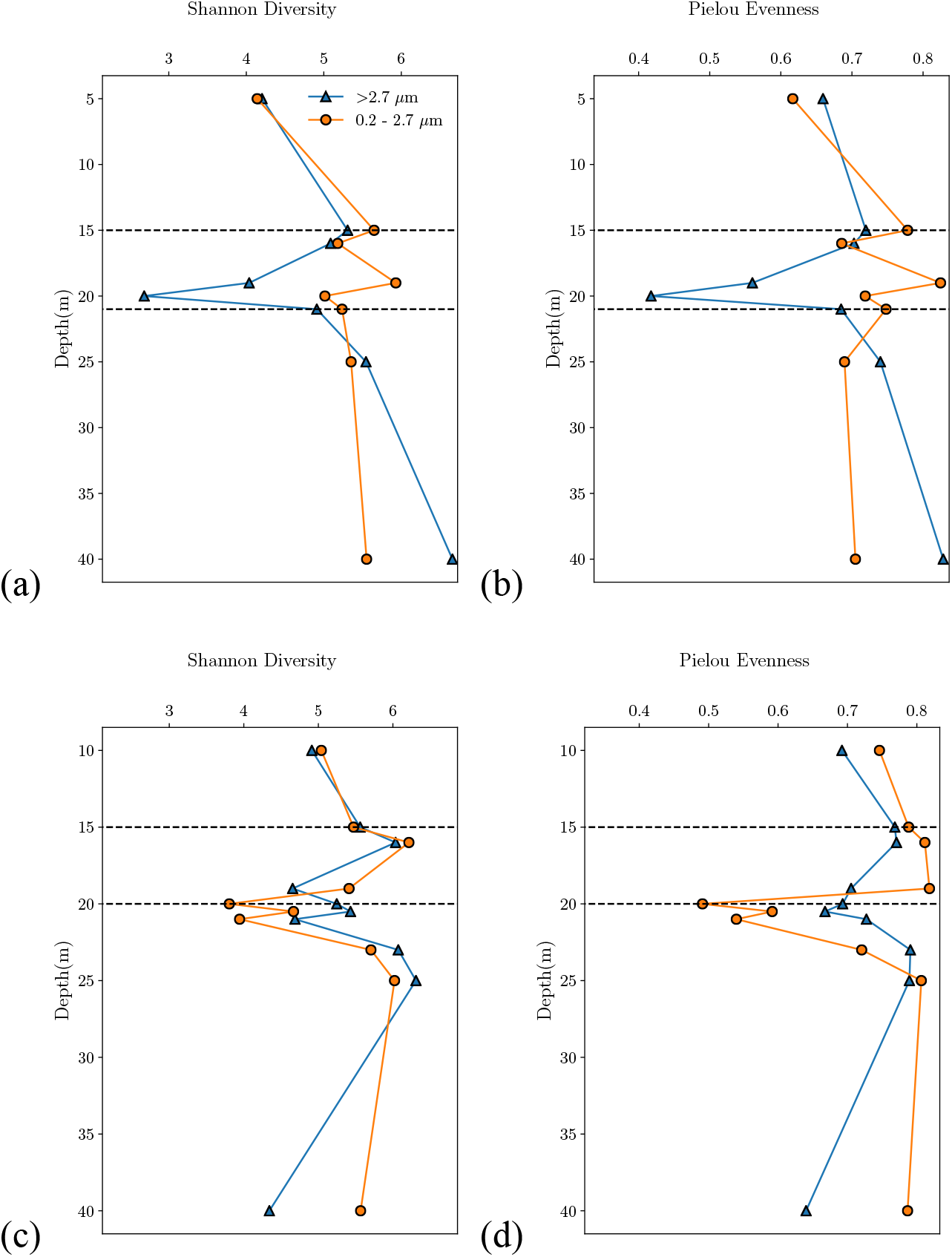
Alpha diversity of PA and FL assemblages as measured by Shannon diversity, *H* (a, c) and Pielou evenness indices (b, d) for Oct (a, b) and Jul (c, d) samples. Broken lines represent oxycline boundaries on those dates.

Excluding the light-scattering layer, alpha diversity generally increased with depth. During both sampling times, FL assemblages were typically more diverse than PA assemblages except within the mixolimnion, where PA and FL assemblages were approximately equally diverse (H = 4-5). One exception was at 40 m during October (PA H=6.8, FL H=5.6). The difference in diversity between PA and FL assemblages was smaller during July than during October (typically by less than 0.5) and remained similar over depth with the exception of the deepest (40 m) sample (PA H=4.3, FL H=5.6). In contrast, during October, differences ranged from 1.5-to 2-fold within the oxycline and was nearly 3-fold within the light-scattering layer. These results suggest that assemblage partitioning varies seasonally. Possibly, this can be attributed to greater rates of physical particle aggregation and disaggregation (and therefore greater similarity) in summer, when particulate matter concentrations are likely higher as a result of the blooms.

To characterize compositional differences among samples (beta diversity), we calculated the Bray-Curtis dissimilarity metric and performed a PCoA on the resulting dissimilarity matrix. The first three coordinates of the Bray-Curtis PCoA explained 53.2% of total sample variance. When metadata variables were mapped over the samples in coordinate space, we found that the FGL community was strongly partitioned by ORP, assemblage-type, and by field campaign (time-of-year). We verified that these community composition groups were significantly different from one another by multiple statistical tests (ANOSIM and PERMANOVA, *p* < 0.05). Detailed Bray-Curtis PCoA PERMANOVA results are provided in supplemental materials (**Supplementary materials** Table S2). ORP separated clusters from one another along PC1 in PC1-PC2 space, accounting for the largest variance (Fig. 5). Interestingly, samples from non-euxinic depths did not further separate from one another along PC1 by redox condition (i.e., suboxic, hypoxic, normoxic). However, samples did separate into two subclusters by sampling date along PC2, primarily due to variations in seasonal surface processes (Fig. 5). Assemblage type (PA or FL) separated clusters from one another along PC3 in PC2-PC3 space (Fig. 5).

**Figure 5.**
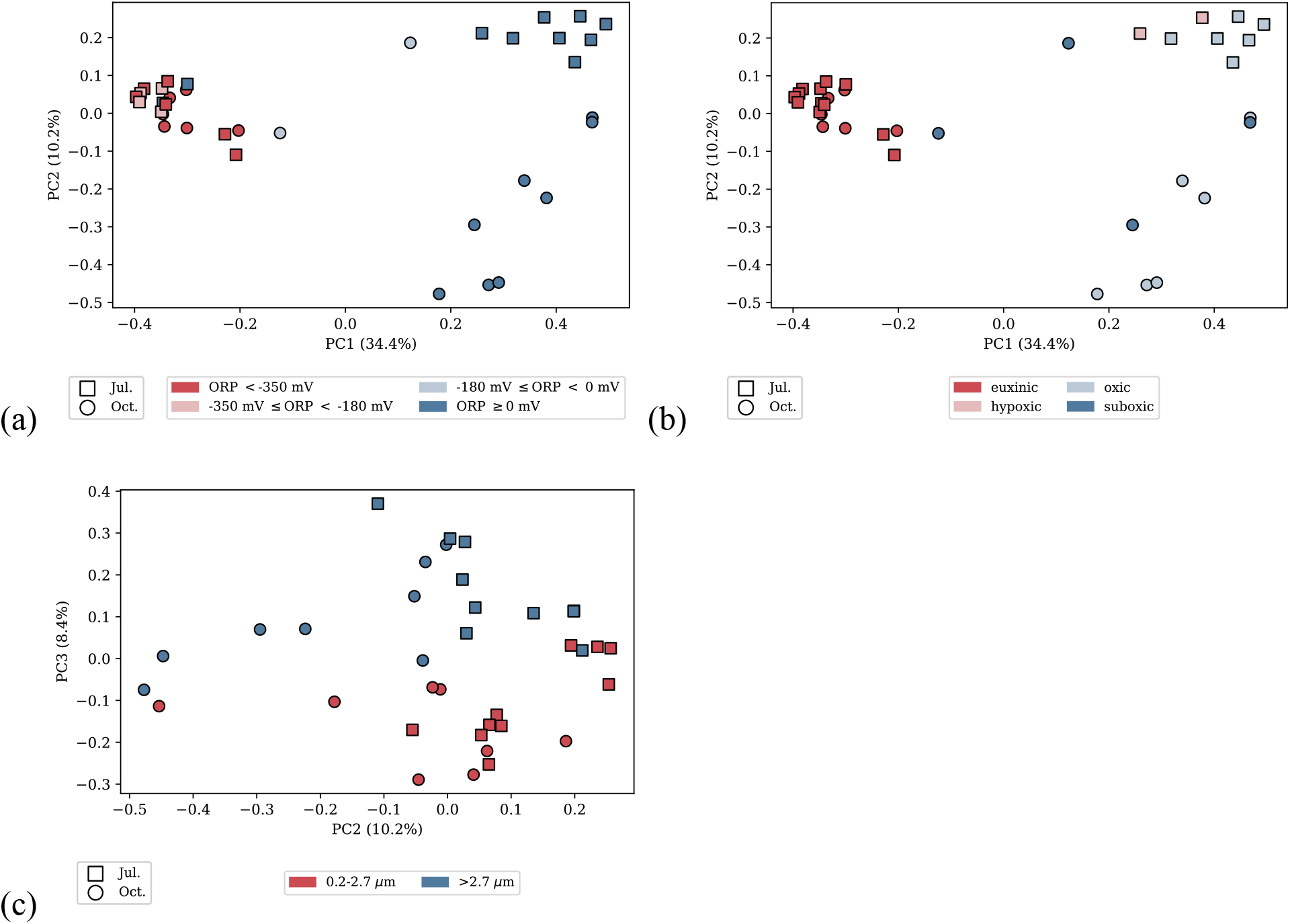
Beta diversity Bray-Curtis PCoA for all samples shown in (a, b) in PC1-PC2 space and (c) PC2-PC3 space. Samples in (a) are color-coded by ORP ranges. Samples in (b) are color coded by redox condition as defined by oxygen concentration and presence or absence of sulfide. Samples in (c) are color-coded as PA (>2.7 *µ*m) and FL (0.2-2.7 *µ*m). Percentages of total variance explained appear on axes.

### Taxonomic composition

We found that the archaeal and bacterial 16S rRNA gene amplicon community composition of the FGL water column during July and October varies with assemblage type (PA or FL), redox-zonation, and time-of-year. Across all depths, Bacteroidetes (Jul.=52.3%, Oct.=26.1%), Proteobacteria (Jul.=21.1%, Oct.=37.3%), Cyanobacteria (Jul.=11.0%, Oct.=11.9%), Actinobacteria (Jul.=6.3%, Oct.=11.0%), and Verrucomicrobia (Jul.=0.9%, Oct.=0.8%) were the most abundant phyla among total reads during both sampling dates (Fig. 6). Amplicons from Bacteroidetes had no discernable depth dependent patterns. Actinobacteria, Cyanobacteria and Verrucomicrobia sequences were more abundant in the mixolimnion and oxycline than in deeper waters by approximately 1 to 1.5 orders of magnitude. The relative abundance of Proteobacteria in 16S rRNA gene libraries decreased over depth from 85.0 to 17.7% between 5 and 40 m in FL assemblages during October. During July, Proteobacteria were far more abundant in the deepest sample (40 m) in both assemblage-types (PA=71.4%, FL=55.8%) than throughout the rest of the water column (PA=4.9-31.7%, FL=11.0-26.4%). During October, however, they had much greater relative abundances in the shallowest sample (5 m) in both assemblage-types (PA=94.4%, FL=85.0%) than throughout the rest of the water column (PA=17.7-61.2%, FL=17.3-51.1%). Chlorophyte algal chloroplast 16S rRNA amplicon sequences were also highly abundant in October, especially near the lower oxycline boundary. Chloroplast depth distriubtions aligned well with those of the chlorophyll-a fluorescence (**Supplementary materials** Figs. S9, S10). Chloroplast sequences were scarce in all July samples.

**Figure 6.**
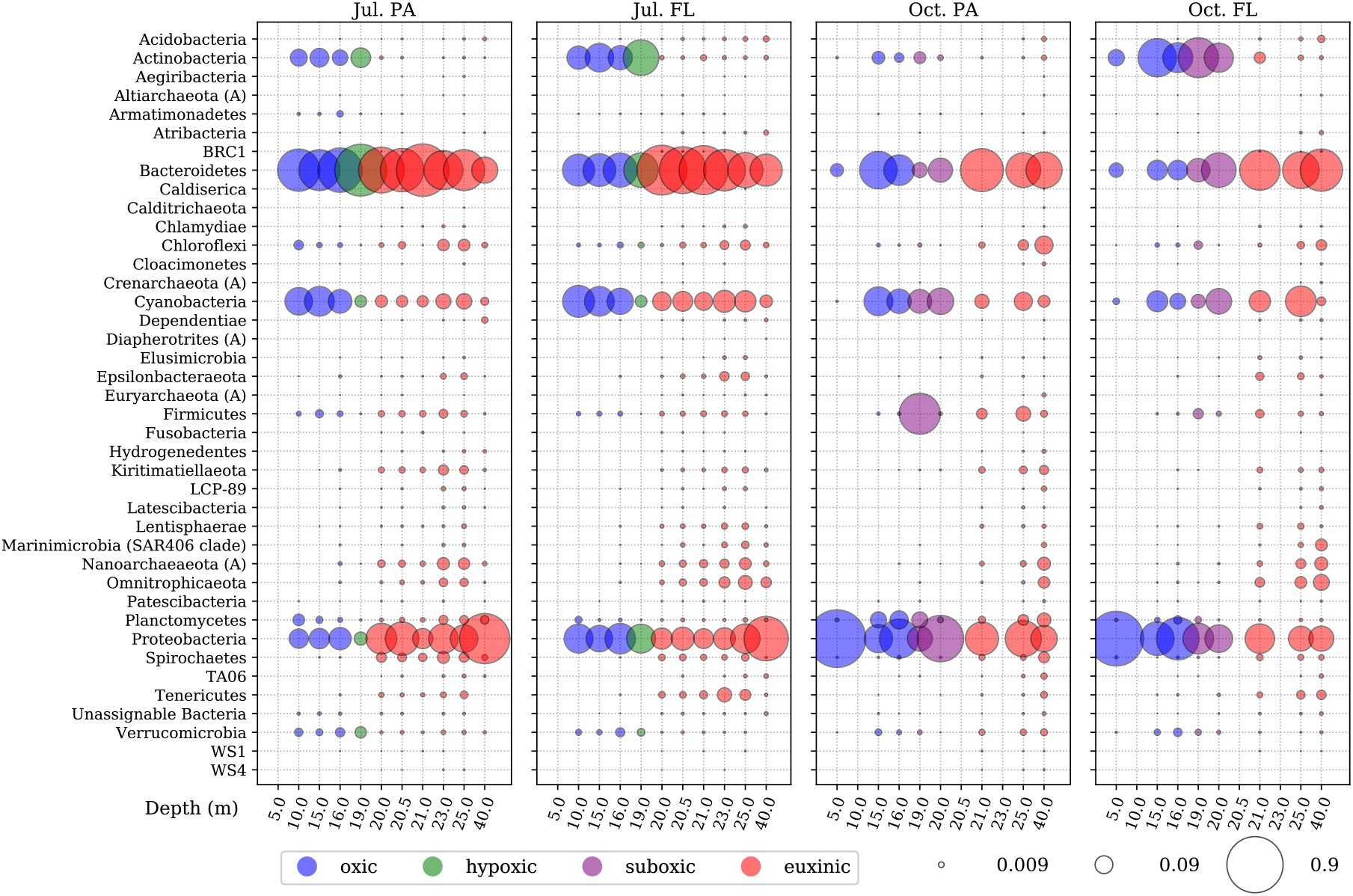
Relative abundances of all recovered phyla in PA and FL assemblages throughout the FGL water column during July 2017and October 2016. Bubble size scales with relative abundance and bubble color corresponds to redox condition.

The most abundant families in the 16S rRNA amplicon libraries varied more strongly with depth than the most abundant phyla (Fig. 7). Family abundances differed most between sampling dates in mixolimnion and oxycline samples, while compositions between sampling dates in the monimolimnion were similar. The most abundant families in the July mixolimnion and oxycline belonged to the Flavobacteriales (PA=23.7-62.7%, FL=9.5-18.0%), Cytophagales (PA=0.7-3.7%, FL=3.2-9.5%), Synechococcales (PA=3.9-24.9%, FL=3.8-28.8%), Frankiales (PA=2.8-6.1%, FL=11.5-31.6%), Microtrichales (PA=2.0-4.0%, FL=2.1-4.9%) and SAR11(PA=0.4-1.8%, FL=2.5-4.0%) orders.

**Figure 7.**
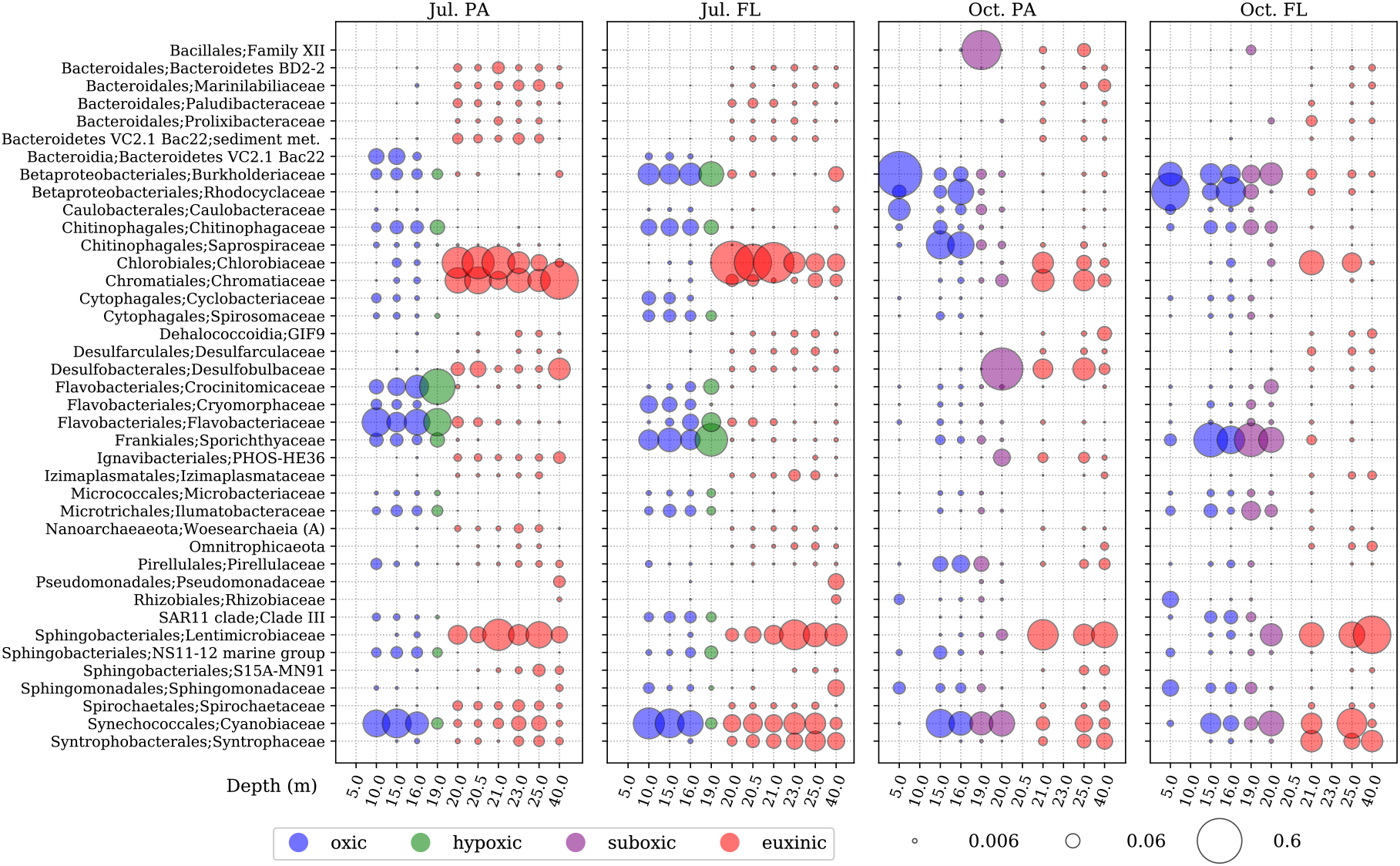
Relative abundances of top families in PA and FL assemblages recovered throughout the FGL water column during July 2017 and October 2016. Bubble size scales with relative abundance and bubble color corresponds to redox condition.

Families belonging to Synechococcales (PA=0.2-23.6%, FL=1.2-18.4%) and Frankiales (PA=0.1-2.7%, FL=4.2-34.5%) orders remained abundant in October, but other families abundant in July represented much less of the October mixolimnion and oxycline assemblages. Instead, families in the Bacillales (PA=0.0-46.1%, FL= 0.0-2.7%), Ignavibacteriales (PA=0.0-8.5%, FL=0.0-0.2%), Betaproteobacteriales (PA=1.4-67.0%, FL=16.9-59.3%), Chitinophagales (PA=1.8-28.7%, FL=1.5-6.5%), Rhizobiales (PA=0.4-4.4%, FL=0.1-10.0%) and Pirellulales (PA=0.2-8.9%, FL=0.0-1.7%) orders were more highly represented. The Caulobacterales and Bacillales orders were represented by single genera (*Brevundimonas* and *Exiguobacterium*, respectively). ASVs from the Burkholderiaceae family (Betaproteobacteria) contributed the greatest number of identifiable genera as well as unclassified members (e.g., *Limnohabitans*, *Polynucleobacter*, *Aquabacterium*, *Hydrogenophaga*, *Ideonella*, *Rhodoferax*). *Ferribacterium* was the sole representative of the Rhodocyclaceae family (Betaproteobacteriales). *Ferribacterium* was by far the most abundant genus in the October mixolimnion and oxycline and was significantly more abundant in October than in July samples (ANCOM test, W-statistic=888).

Consistent with other field observations (see **Section 4.1**), the most abundant families in the light-scattering layer (19-23 m) were photoautotrophic Cyanobiaceae (cyanobacteria), Chromatiaceae (purple sulfur bacteria) and Chlorobiaceae (green sulfur bacteria). In the light-scattering layer, these families combined made up 3.9-54.4% of PA assemblages and 3.9-68.9% of FL assemblages during July and 16.7-33.3% of PA assemblages and 5.5-30.4% of FL assemblages during October. Within this layer, the Desulfobulbaceae and PHOS-HE36 were also especially abundant. Desulfobulbaceae sequences represented a single genus capable of sulfate reduction and sulfur disproportionation, *Desulfocapsa*. Desulfobulbaceae (PA=0.1-52.9%, FL=0.0-0.6%) and PHOS-HE36 (PA=0.0-8.5%, FL=0.0%) sequences were more abundant in October, when photoautotroph abundances were lower. Below the light-scattering layer, the relative abundance of purple sulfur bacteria decreased with depth but was highly enriched in the deepest July sample (40 m, PA=42.5%, FL=4.5%). Chlorobiaceae (green sulfur bacteria) decreased with depth in both July and October by approximately an order of magnitude between their maximum within the light-scattering layer (21 m) and the deepest sample (40 m). Retrieval of photoautotroph sequences from depths below the photic zone likely represents sedimenting senescing or dead cells, indicating that material is being vertically transported from the light-scattering layer towards the lakebed. Depth distributions and the assemblage-partitioning of Desulfobulbaceae and purple sulfur bacteria sequences were similar, suggesting physical association in particles.

The most abundant taxa recovered from below the light-scattering layer during both field campaigns were organotrophic anaerobes. These included sequences from the sulfate-reducing families Desulfarculaceae (consisting entirely of *Desulfatiglans*) and Syntrophaceae (combined: Jul. PA=1.8-3.4%, FL=8.9-12.4%; Oct. PA=5.7-7.7%, FL=7.9-14.4%). Fermentative and/or respiratory organotrophic taxa were highly represented by ASVs from the Lentimicrobiaceae, Spirochaetaceae (entirely *Sphaerochaeta*), Prolixibacteraceae, Marinilabiliaceae, and families belonging to the order Bacteroidales (Bacteroidetes BD2-2, VC2.1 Bac22). Below the light-scattering layer these families combined accounted for 11.0-32.3% of PA assemblages and 16.4-21.7% of FL assemblages during July and 17.7-30.0% of PA assemblages and 23.4-46.9% of FL assemblages during October. Several putative organohalide-respiring taxa were also abundant, including the environmental family S15A-MN91 and the GIF9 order of Dehalococcodia (Jul. PA=2.3-5.6%, FL=1.0-2.8%; Oct. PA=3.3-9.0%, FL=1.4-3.2%). An unassignable member of the archaeal class Woesearchaeia was also abundant at one depth (25 m) during July (PA=1.7%, FL=1.0%) (Fig. 7).

### Assemblage partitioning

The assemblage-partitioning (PA or FL) of archaeal and bacterial 16S rRNA gene amplicons was evident throughout the FGL water column during July and October at the family level. Of the five most abundant phyla, only Actinobacteria were more abundant in one type of assemblage (FL). As expected, algal chloroplasts were primarily captured in PA assemblages because they represent phototrophic eukaryote cells. Assemblage partitioning of some families differed between sampling dates. For example, families in the Flavobacteriales order were highly over-represented in PA assemblages throughout the water column in July, but neutral or even slightly over-represented in FL assemblages in October (Fig. 8). In contrast, families that were most abundant in the light-scattering layer, such as Chromatiaceae, Desulfobulbaceae, Pirellulaceae and PHOS-HE36, were recovered almost entirely from PA assemblages during both campaigns. An ASV classified as the purple sulfur bacterium, *Thiodictyon*, was significantly more abundant in PA assemblages than in FL assemblages in all samples (ANCOM test, W-statistic=885).

**Figure 8.**
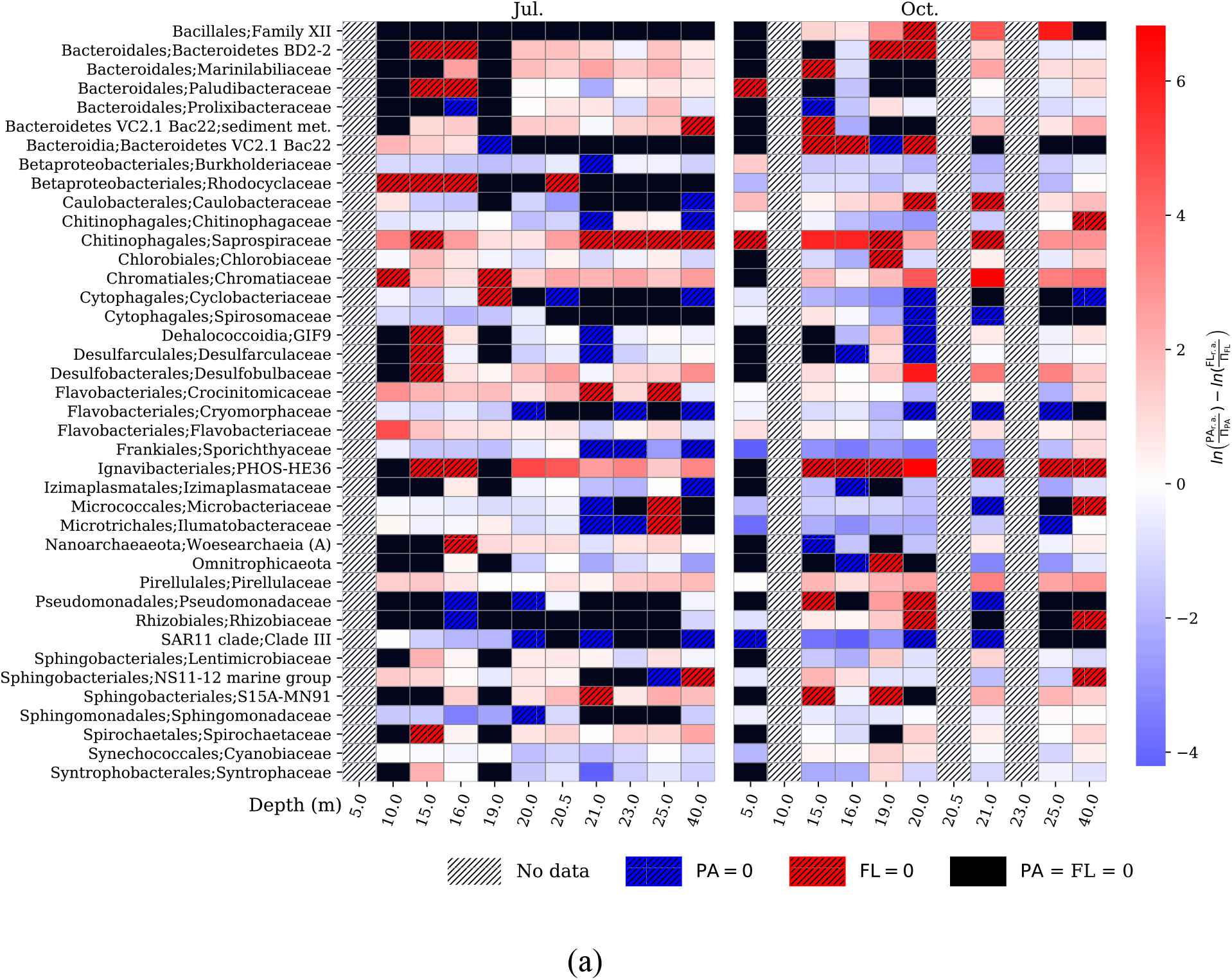
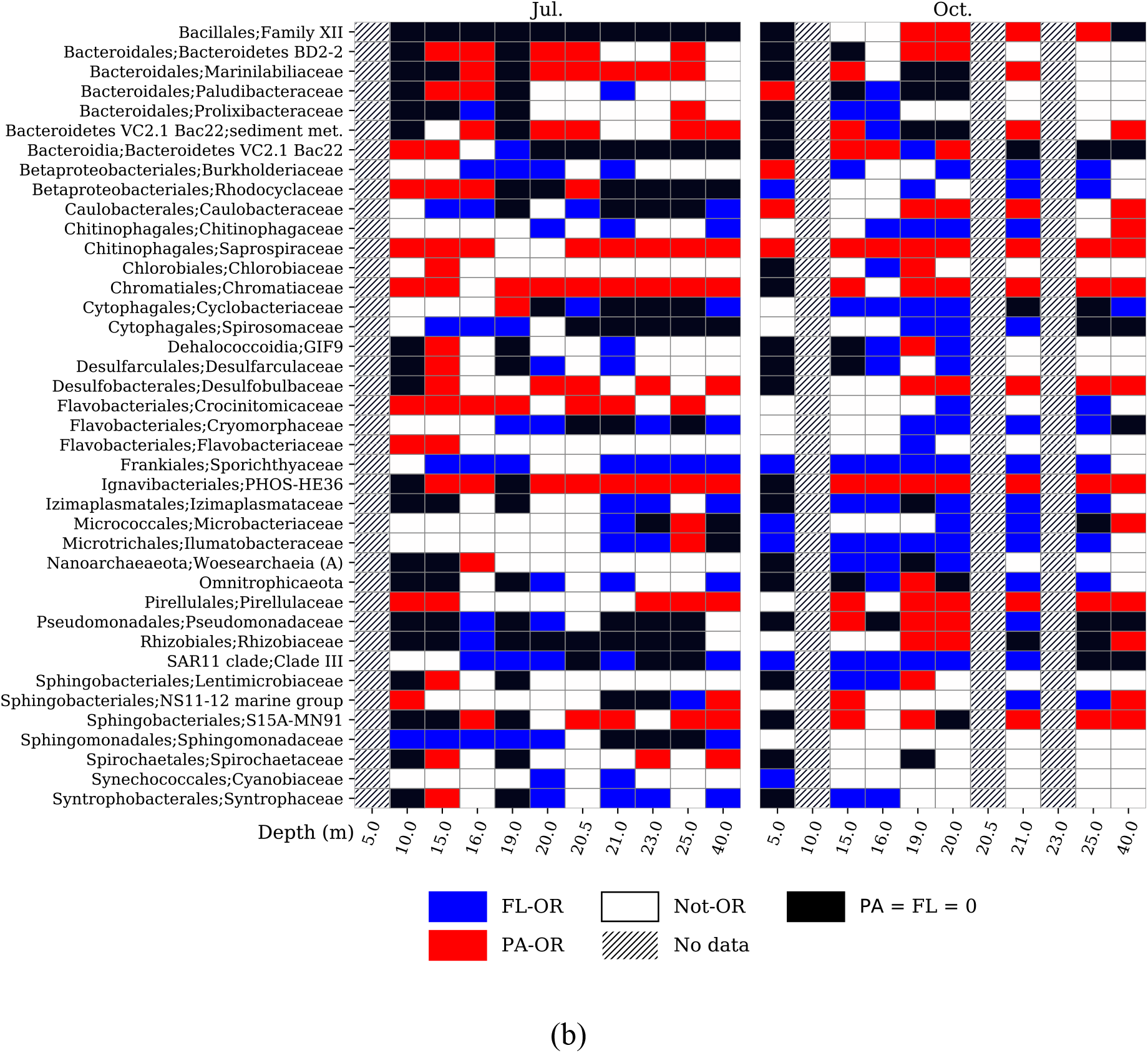
(a) Overrepresentation factors (OFs) of top families recovered throughout the FGL water column during July 2017 and October 2016. Positive (red) OFs indicate overrepresentation in the PA assemblage, while negative (blue) values indicate overrepresentation in the FL assemblage. (b) The same data classified by overrepresentation factor as over-represented in particle-associated assemblages (≥ 1.25, PA-OR), over-represented in free-living assemblages (≤ −1.25, FL-OR), or neither (Not-OR).

Sulfate-reducing bacteria, organohalide-respirers, and other fermentative and/or respiratory organotrophic families that were highly abundant below the light-scattering layer were low-abundance, but highly over-represented (over-representation factor ≥ 2, or ≥ 7.4-fold more abundant) in PA assemblages within the mixolimnion and shallow oxycline (Fig. 8). More taxa, and notably those that are less oxygen-tolerant, displayed this pattern in July, possibly due to enhanced thermal stratification in the upper water column. These included Spirochaetaceae, Desulfobulbaceae, Desulfarculaceae, Syntrophaceae, and Bacteroidetes BD2-2 during July, GIF9, S15A-MN91 and VC2.1 Bac22 during October, and Marinilabiliaceae during both field campaigns.

Because the most abundant families belong to four phyla and have only a few putative metabolisms, over-representation factors of all phyla were considered to look for less abundant, strongly partitioned, and potentially functionally important taxa (Fig. 9). During both field campaigns, the most over-represented PA phyla throughout the water column included Planctomycetes, Firmicutes, and Verrucomicrobia. Overrepresentation was greatest (OF ≥ 2, or ≥ 7.4-fold more abundant) at the shallowest and deepest sampling depths, suggesting they originate from the mixolimnion and are vertically transported in particles to the deep monimolimnion. Supporting this hypothesis, members of these phyla often attach to sinking particles in the shallow waters and therefore have this type of over-representation distribution (Delong et al., 1993, Bižić-Ionescu et al., 2014a, Thiele et al., 2015, Duret et al., 2018). Anaerobic phyla that were moderately abundant in the monimolimnion were rare but highly over-represented in PA assemblages in the July mixolimnion and oxycline samples, such as Marinimicrobia (SAR406), Atribacteria (entirely the class JS1), and Nanoarchaeota.

**Figure 9.**
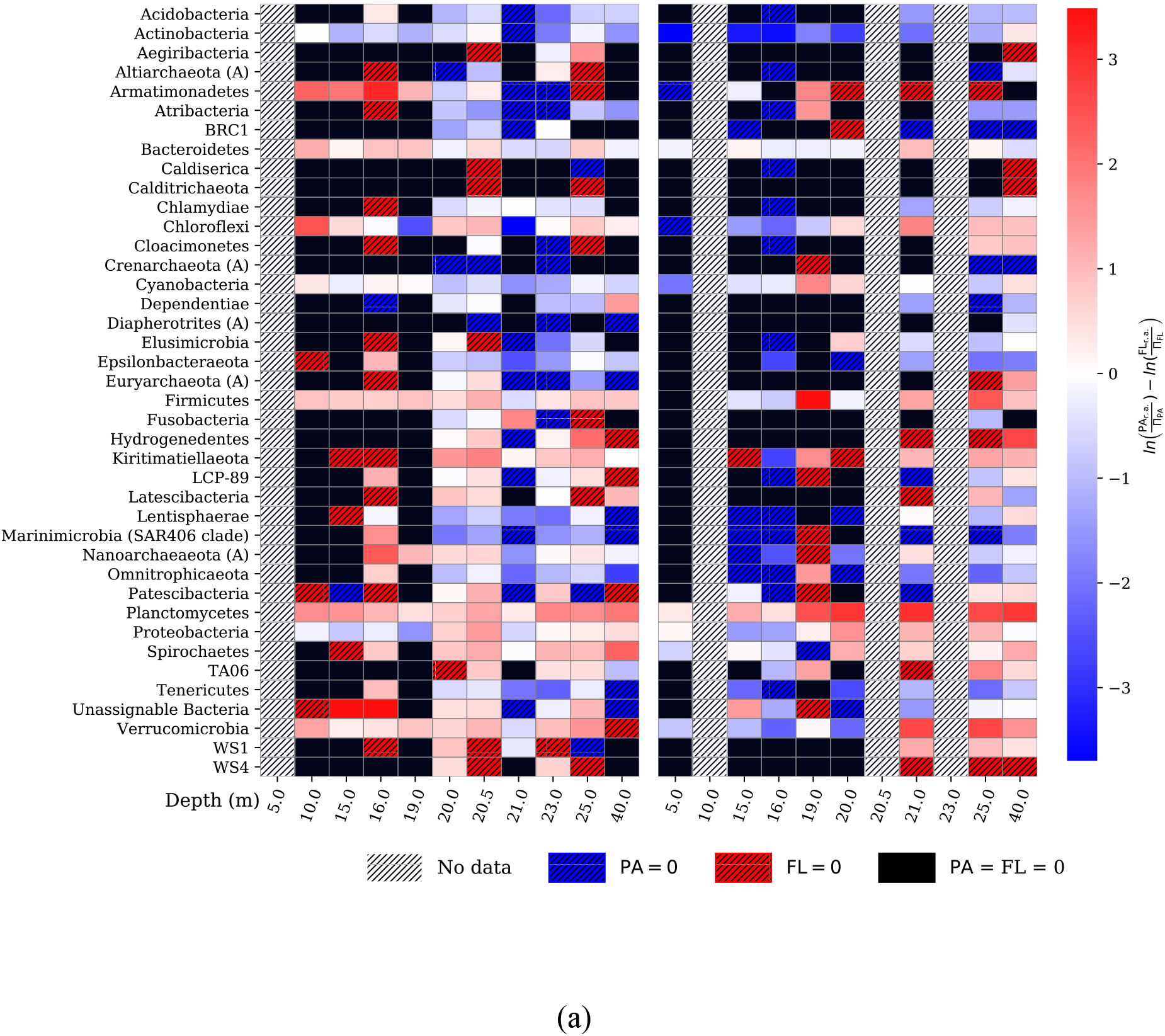
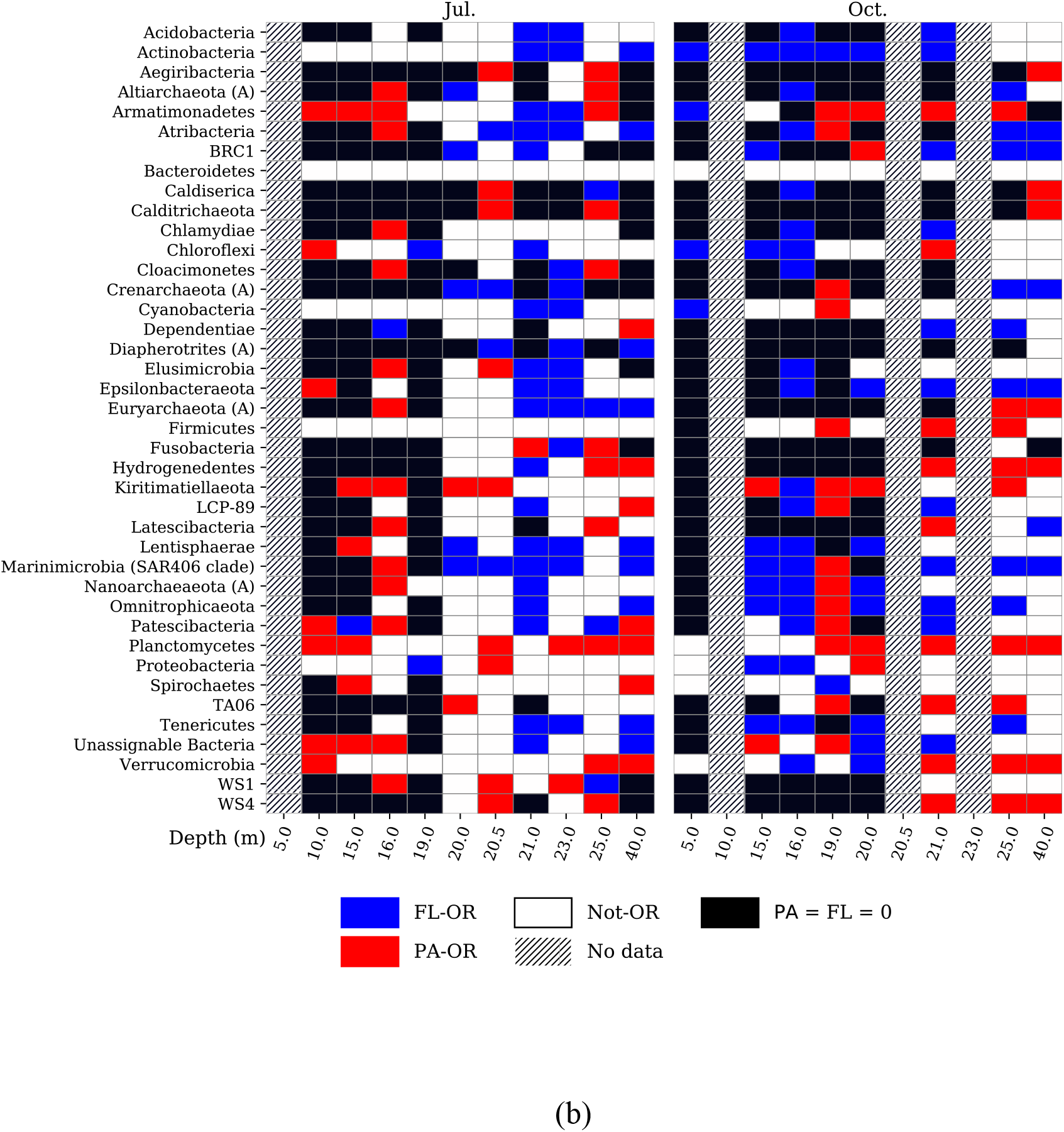
(a) Overrepresentation factors (OFs) of all recovered phyla throughout the FGL water column during July 2017 and October 2016. Positive (red) OFs indicate overrepresentation in the PA assemblage, while negative (blue) values indicate overrepresentation in the FL assemblage. (b) The same data classified by overrepresentation factor as over-represented in particle-associated assemblages (≥ 1.25, PA-OR), over-represented in free-living assemblages (≤ −1.25, FL-OR), or neither (Not-OR).

Most over-represented phyla in PA assemblages were low abundance anaerobes and often represented by one or two families. These included Altiarchaeota, Armatimonadetes, Chloroflexi, Euryarcheaota, Hydrogenedentiales, Kiritimatiellaeota and Spirochaetes. Euryarcheota sequences consisted entirely of an uncultured methanogenic order Methanofastiodiosales and the methanogen *Methanoregula*. Altiarchaeota sequences arose entirely from the uncultured class Altiarchaeia. Hydrogenedentiales was entirely represented by the family Hydrogenedensaceae. Spirochaetes sequences were entirely from the family Spirochaetaceae.

## Discussion

### Particles shape microbial composition and diversity

Microbes associated with particles are taxonomically and functionally distinct from free-living microbes in aquatic environments, but are intertwined with one another in biogeochemical interplay. Differences are shaped by the availability of organic substrates, nutrients, reductants, and oxidants required for microbial metabolisms. However, their interactions are driven by cooperation, antagonism, symbiosis and syntrophy and the physical exchange of dissolved and particulate reactants. In permanently redox-stratified water bodies, these assemblages are additionally shaped by ambient redox conditions, which determines the availability of reductants and oxidants over depth. Lakes play a critical role in global carbon sequestration (Downing et al., 2008), and many are likely to become permanently redox-stratified due to global climate change (Lau et al., 2020). However, studies of PA and FL microbial assemblages in permanently redox-stratified water bodies have primarily focused on larger marine systems (Fuchsman et al., 2011, Suter et al., 2018, Suomenin et al., 2020).

This study revealed that the community phylogenetic composition of all samples from FGL varied among redox condition (oxidation reduction potential) and assemblage type (PA or FL), while only samples from non-sulfidic depths differed by sampling date (Fig. 6). In permanently stratified water columns, deeper, anoxic waters are usually isolated from surface seasonal physico-chemical changes by a pycnocline, a water density gradient. Therefore, the nearly temporally invariant geochemistry at depth likely also shapes community composition independently of seasonal variables (e.g., grazing pressure, wind mixing, organic matter production and flux, temperature).

Richness and evenness of the microbial assemblages also varied with redox condition and assemblage type, with differences between assemblages varying temporally. Consistent with other studies of redox-stratified water bodies and of FGL, alpha diversity generally increased over depth, except within the light-scattering layer (Fuchsman et al., 2011, Suter et al., 2018, Suomenin et al., 2020, Rojas et al., 2021, Block et al., 2021). However, alpha diversity was greater in FL assemblages than in PA assemblages, contrary to what has been reported in marine anoxic basins (Fig. 5a,b) (Fuchsman et al., 2011, Suter et al., 2018, Suomenin et al., 2020). In FGL, this may reflect vertical transport of low-diversity PA assemblages from the mixolimnion and light-scattering layer towards the lakebed by gravitational settling. This hypothesis is supported by the presence of highly over-represented taxa from the mixolimnion and light-scattering layer at depth (Figs. 6, 7, 8, 9). Additionally, recalcitrant particulate organic substrates that have been vertically-transported to the monimolimnion (e.g., chitin, lignin, cellulose) may select for specialist assemblages of lower diversity (Rojas et al., 2021).

### Composition of the mixolimnion and oxycline differs by sampling date

In aquatic environments, nutrient and organic matter availability varies seasonally, resulting from a combination of physical mixing, primary production, and other biological activity (Sarmiento and Gruber, 2013). In FGL, as in all permanently redox-stratified water bodies, these processes are restricted to water above the pycnocline, unless there are lateral oxygen intrusions. Compositional differences between the July and October mixolimnion and oxycline assemblages in FGL likely reflect seasonal availability of organic carbon substrates and inorganic nitrogen. The concentration and quality of organic carbon substrates is likely lower in October than in July because the shallow cyanobacteria bloom (a major source of labile dissolved and particulate organic carbon) had ended. At the same time, inorganic nitrogen availability is higher in October than in July (Fulton et al., 2018), likely due to lower uptake by the senescing shallow cyanobacteria population and deeper wind-mixing (Fig. 10).

**Figure 10.**
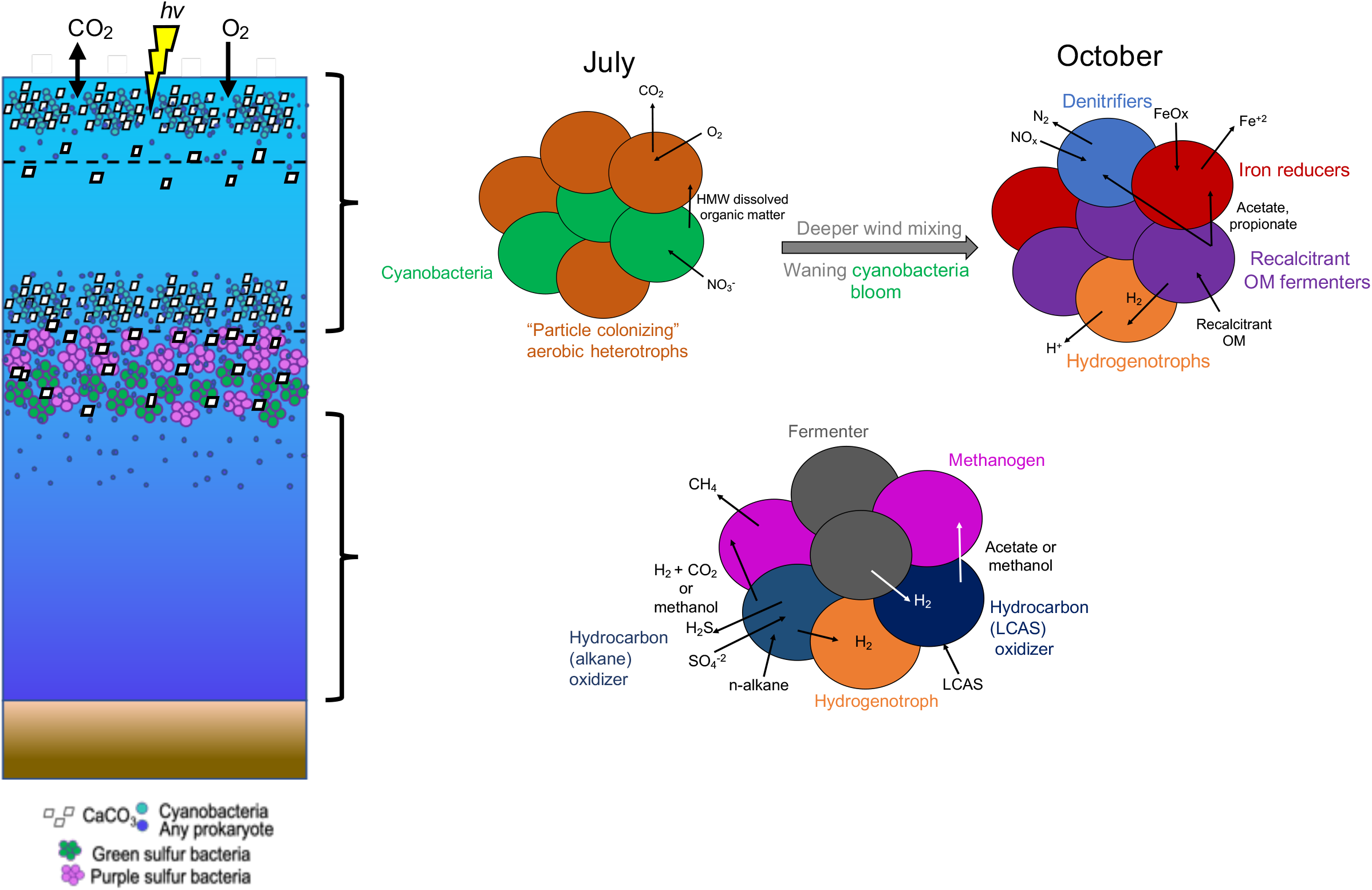
A conceptual diagram of major microbial processes in the FGL water column. Cyanobacteria populations in the mixolimnion and near the lower oxycline boundary carry out oxygenic photosynthesis and cause CaCO_3_ precipitation. Green and purple sulfur bacteria at the lower oxycline boundary and in the upper monimolimnion mediate photoautotrophic sulfur oxidation. In July, the mixolimnion is dominated by PA aerobic heterotrophs that colonize algal biomass. In October, the mixolimnion and oxycline instead hosts abundant denitrifiers, iron reducers, hydrogenotrophs, and fermenters. Below the light-scattering layer, the seasonally invariant monimolimnion community consists of taxa that are common in hydrocarbon oxidizing, methanotrophic, syntrophic consortia. HMW=High molecular weight. LCAS=Long chain alkyl substituted

We hypothesize that lower organic C to inorganic N ratios in October should favor nitrogen removal from FGL through denitrification and anaerobic oxidation of ammonia (anammox) over nitrogen recycling by the dissimilatory reduction of nitrate to ammonium (DRNA) (Nizzoli et al., 2010; Chutivisut et al., 2018). Supporting this hypothesis, dissolved organic carbon concentrations in FGL are nearly invariant between summer and autumn (Havig et al., 2015, 2018), while nitrate concentrations are nearly two-fold greater in autumn than summer. As a result, the dissolved organic C to inorganic N ratio is nearly two-fold lower in autumn. Therefore, the taxonomic composition of the July mixolimnion and oxycline should reflect organic matter abundance (aerobic heterotrophy) and inorganic nitrogen scarcity (diazotrophy), while the composition of the October mixolimnion and oxycline should reflect nitrogen-removal (denitrification, anammox) and organic matter scarcity (oxidation and/or fermentation of recalcitrant organic carbon substrates). Indeed, the most abundant families in the July mixolimnion and oxycline were aerobic heterotrophs and diazotrophs, while the most abundant families in the October mixolimnion and oxycline were putative and known denitrifiers, iron-reducers, and fermenters. Although the abundance of the obligatory anammox bacterium *Candidatus anammoximicrobium* in the oxycline did not vary between July and October, the taxon’s assemblage parititioning did; it was almost entirely recovered from PA assemblages in the July oxycline and almost entirely recovered from FL assemblages in the October oxycline.

#### July mixolimnion and oxycline

The Crocinitomicaceae and Flavobacteriaceae families within Flavobacteriales order were especially abundant and over-represented in July PA assemblages (Figs. 8, 9, and 10). These families metabolize complex, proteinaceous substrates that other microbes are unable to access directly. Therefore, they are dominant during algal blooms in freshwater and marine environments, and are integral to initiating particle remineralization (DeLong et al., 1993, Bižić-Ionescu et al., 2014a, b, Duret et al., 2019). The algicidal Cytophagales order, which is often associated with particles and functionally similar to the Flavobacteria (Kirchman, 2002), were more abundant in FL assemblages (Figs. 8, 9). However, there are reports of the Cytophagales families being most abundant in FL assemblages during freshwater cyanobacteria blooms (Akins et al., 2018).

Consistent with other niche partitioning studies during freshwater algal blooms, the other abundant aerobic heterotrophs (families within SAR11 and Microtrichales) and diazotrophs (families within Frankiales) were over-represented in FL assemblages (Fig. 9) (Salcher et al., 2011, Bižić-Ionescu et al., 2014b, Morrison et al., 2017, Akins et al., 2018). Diazotrophy is advantageous under the nitrogen-limited conditions typical of algal blooms (Shin et al., 2019) because nitrogen requirements can be met with little available nitrate. These taxa do not access their organic substrates within or on particles like the Flavobacteria. However, they are associated with blooms because the labile dissolved organic substrates they consume originate from particles and algal lysates, in part, from the activity of microbes like the Flavobacteria (Mayali and Azam, 2004, Alonso and Pernthaler, 2006, Salcher et al., 2011, Zheng et al., 2020). The interconnectedness of seasonal particle production (biomass) and consumption of labile dissolved organic substrates is hinted at by the overlapping distributions bacterial heterotrophic production and phycoerythrin concentrations in the July profiles. This alignment suggests that bacterial heterotrophic production is responsive to cyanobacteria abundances (Figs. 2c, 3a).

#### October mixolimnion and oxycline

The most abundant families in the October mixolimnion and oxycline appear to have a competitive advantage when organic carbon substrates are scarce and less labile and/or when inorganic nitrogen is more abundant. Under these limitations, the putative metabolisms of the taxa over-represented in PA assemblages (Pirellulaceae, families within Chitinophagales, Rhizobaceae and Burkholderiaceae, Figs. 8, 9), are likely connecting multiple elemental cycles (iron, nitrogen, hydrogen, and carbon) in particles. For example, many of the Burkholderiaceae genera, including *Aquabacterium, Hydrogenophaga*, *Ideonella* and unclassified Burkholderiaceae, are capable of consuming aromatic compounds or hydrocarbons (e.g., pyrene, benzene and petroleum hydrocarbons) that are integral to producing volatile fatty acids for co-existing iron-reducers (Kalmbach et al., 1999, van der Zaan et al., 2012, Zhang et al., 2016, Yan et al., 2017, Wieczorek et al., 2019, Liu et al., 2020, Fig. 10).

Burkholderiaceae-rich consortia have also recently been linked to enhanced denitrification, which could be attributable to the metabolic flexibility of the most common genera. Metabolic flexibility to mediate organotrophic and/or chemoautotrophic nitrogen and iron cycling reactions could be beneficial when organic substrates are scarce. (Zhang et al., 2016, Liu et al., 2020). For example, *Hydrogenophaga* can couple hydrogen oxidation with denitrification (Liu et al., 2020) or aromatic carbon oxidation with iron reduction (Yan et al., 2017). *Aquabacterium* can mediate iron oxidation or chemoorganotrophic denitrification (Kalmbach et al., 1999, Zhang et al., 2016). Notably, these were also the most common identifiable genera of Burkholderiaceae during October.

A potential connection between recalcitrant organic carbon oxidation and iron reduction in FGL during October is supported by the much higher abundance of obligate iron-reducers in the mixolimnion and oxycline compared to July, including *Ferribacterium* (Rhodocyclaceae, Figs. 8, 9) and *Rhodoferax* (Cummings et al., 1999, Finneran et al., 2003, Chan et al., 2018). Iron-reducers were expected to be more common in PA assemblages, because reducible iron is particulate. However, *Ferribacterium* and *Rhodoferax* sequences were equally likely to be recovered in both assemblage types (over-representation factor ∼ 0). This unexpected assemblage partitioning of iron-reducers could be explained by the documented existence of particulate-iron-rich extracellular polymeric substances (EPS) at non-sulfidic depths in FGL (Kamennaya et al., 2020), as iron cycling is promoted in EPS (Chiu et al., 2017). If iron-reducers do reside in EPS in FGL, which is easily disturbed during filtration (Engel, 2009), they could pass through the 2.7 *µ*m filter.

The high abundance of the above-described taxa and other denitrifiers that are dominant in natural waters and bioreactors (e.g., Pirellulaceae, PHOS-HE36, *Exiguobacterium*), especially those with low organic C to inorganic N ratios, points to overall nitrogen-removal from FGL in October (Dabert et al., 2001, Singh et al., 2015, Kelogg et al., 2016, Gutiérrez-Preciado et al., 2017, Zhou et al., 2019). Other than the anaerobic ammonium oxidizer, *Candidatus anammoximicrobium*, taxa within the Pirellulaceae family exclusively mediate nitrification and organotrophic denitrification (Kelogg et al., 2016, Song and Liu, 2019). Over-representation of these taxa in PA assemblages suggests that particles are important microhabitats for denitrifiers under subsaturated oxygenated conditions as found in other studies (Fuchsman et al., 2011, Ganesh et al., 2015, Suter et al., 2018).

### Anaerobic remineralization in the monimolimnion

In permanently redox-stratified water bodies, deeper, anoxic waters host diverse assemblages of anaerobes that derive energy from fermentation, chemoorganotrophic sulfate, iron and manganese oxide reduction, chemoautotrophic oxidation or disproportionation of reduced sulfur species, and the chemoautotrophic oxidation of iron, manganese, ammonium, and methane. Due to low energy yields and the need to remove inhibitory metabolic byproducts (e.g., formate and hydrogen during fermentation), microbes mediating these reactions are often physically associated in cellular aggregates (Sieber et al., 2012, Morris et al., 2013). Physical association facilitates the metabolic cooperation of microbes that mediate different reactions because nutrients, oxidants, reductants, and electrons can be easily exchanged. This type of cooperation is especially common in energy limited, low-O_2_ environments because metabolic reactions that alone yield too little energy to sponateously occur (endergonic) can become energetically favorable (exergonic) when combined and benefit all participants (=syntrophy). As such, multiple parts of elemental cycles (e.g., sulfate reduction and sulfide oxidation, iron oxidation and reduction) and different elemental cycles (methanogenesis and sulfate reduction) have synergistic interplay in particles.

During October and July, excluding taxa originating from the light-scattering layer and above (likely dead cells), the most abundant families in the monimolimnion were similar. These taxa broadly fall into two categories. Those in the first category are capable of metabolizing complex organic carbon substrates (e.g., carbohydrates, chitin, cellulose, lignin), and likely play a role in initiating anaerobic remineralization of biogenic or autochthonous debris that have been vertically transported towards the lakebed by gravitational settling. Those in the second category are common in methanogenic, sulfate-reducing, hydrocarbon-oxidizing consortia that are characteristic of petroleum-rich environments (Fig. 10). In petroleum-free environments, hydrocarbons can instead originate from decomposing primary producer cells, especially cyanobacteria (Lea-Smith et al., 2015, Love et al., 2021). If this is the case in FGL, the anaerobic remineralization of export primary production in the deep monimolimnion could contribute to hydrogen sulfide, methane, and low molecular weight dissolved organic matter production through hydrocarbon oxidation. Within both categories, further niche-partitioning appears to be driven by putative organic carbon substrate usage (e.g., alkanes versus aromatic compounds, proteins versus carbohydrates).

The families that likely play a role in initiating the remineralization of biogenic or autochthonous debris below the light-scattering layer include Lentimicrobiaceae, Marinilabiliaceae, families in the Bacteroidales order (Bacteroidetes BD2-2, and Prolixibacteraceae), and the archaeal Woesarchaeia. Lentimicrobiaceae ferment polysaccharides (Sun et al., 2016), a common component of algae and their associated mucopolysaccharides or of plant debris. Marinilabiliaceae, Bacteroidetes BD2-2, Prolixibacteraceae, and the archael class Woesarchaeia are able adhere to particles and remineralize proteins and polysaccharides (Mei et al., 2020, Suominen et al., 2020). Notably, not only are Marinilabiaceae associated with algal blooms, but sometimes exclusively scavenge compounds from decaying cyanobacteria (Smith et al., 2017, Ben Hania et al., 2017). This hints that some of the decaying biomass being turned over in the monimolimnion is cyanobacterial.

The families that are common in methanogenic, sulfate-reducing, hydrocarbon-oxidizing consortia include Desulfarculaceae, Syntrophaceae, VC2.1 Bac22, *Sphaerochaeta*, GIF9, S15-MN91, Bathyarchaeota, Methanofastidiosales, and possibly *Rhodopseudomonas*. The representative Desulfarcularceae sequences in this study arose from a single genus, *Desulfatiglans*, which shares the most similarity (95.7% API) with the genome reference sequence *Desulfatiglans anilini* DSM 4660 (RefSeq no. NZ_AULM01000006.1). This species, belonging to group 1 of *Desulfatiglans*, inhabits sulfate-rich environments, and oxidizes aromatic compounds, organosulfonates, and organohalides by dissimilatory sulfate reduction (Jochum et al., 2018). Notably, many of the group 1 *Desulfatiglans* reference sequences have been recovered from petroleum-rich environments (Jochum et al., 2018). The other sulfate-reducing family, Syntrophaceae, is likewise abundant in crude oil-replete environments. However, they oxidize hydrocarbons instead of more refractory organic carbon compounds (e.g., alkane, xylene, or toluene, with alkanes being the preferred substrate) (Edwards and Grbić-Galić, 1994, Gray et al., 2011). The byproducts of hydrocarbon oxidation are consumed by syntrophic methanogens, linking the carbon and sulfur cycles (Gray et al., 2011).

The obligate and putative organohalide-respirers, GIF9 and S15-MN91, also oxidize hydrocarbons in partnership with methanogens (Nedelkova et al., 2005, Hug et al., 2013). They instead use the more refractory long chain alkyl substituted hydrocarbons that are left behind after the preferential biodegradation of lower molecular weight hydrocarbons like alkanes (Cheng et al., 2019). This suggests even further niche partitioning of dissolved organic carbon consumers by hydrocarbon type. The fermentative taxa, including *Sphaerochaeta* and VC2.1 Bac22, have also been recovered from petroleum reservoirs and methane seeps, and found in physical association with sulfate-reducing bacteria, suggesting that they are also part of hydrocarbon-oxidizing, sulfate-reducing, methanogenic consortia (Trembath-Reichert et al., 2016, Dong et al., 2018, Grouzdev et al., 2018, Kleindienst et al., 2012, 2020).

Methanogen sequences were recovered throughout the monimolimnion, supporting the hypothesis that all these organisms are involved in methane production. These included unclassified Bathyarchaeia (now Bathyarchaeota), Methanofastidiosales, *Methanoregula*, and the putative bacterial methanogen *Rhodopseudomonas* (Zheng et al., 2018). Other than *Rhodopseudomonas*, present only during July in the 40 m FL assemblage (3.6%), methanogens were present at low abundances (<0.05% in any assemblage). Methanogen composition appeared to be structured over depth by ambient geochemical conditions and possibly, competition with sulfate reducing bacteria for substrates (e.g., hydrogen, acetate). The only hydrogenotrophic methanogen (*Methanoregula*, Beaver et al., 2016, 2021), was restricted to 20 and 20.5 m, where hydrogen sulfide concentrations, and presumably rates of microbial sulfate reduction are low. Notably, *Methanoregula* is often associated with particulate iron oxide (Beaver et al., 2016, 2021), as would be expected at these depths due to an active iron redox shuttle (Havig et al., 2015).

In the deeper monimolimnion, where high concentrations of hydrogen sulfide presumably originate from bacterial sulfate reduction, methanogens will be easily outcompeted for acetate and hydrogen by sulfate-reducers (Lovely et al., 1982, Schönheit et al., 1982, Robinson and Tiedje, 1984). The most abundant methanogens were therefore Bathyarchaeota and Methanofastidiosales. Bathyarchaeota can use carbon substrates that most methanogens cannot, such as detrital proteins, polysaccharides, aromatic compounds, and methylated compounds; Methanofastidiosales is a methylotrophic methanogen (Evans et al., 2015, Nobu et al., 2016, Zhou et al., 2018). These methanogens can likely make a living co-existing with sulfate-reducers because they are not restricted to using hydrogen and acetate. Finally, *Rhodopseudomonas* was the dominant putative methanogen only in the deepest monimolimnion sample (40 m). *Rhodopseudomonas* produces CH_4_ by fixing CO_2_ using the iron-only nitrogenase when there is a high dissolved iron to molybdenum ratio and a high CO_2_ concentration (Zheng et al., 2018). These conditions are met in the deep monimolimnion (Havig et al., 2015). Its presence only during July could be explained by the secondary chemocline in the deep lake at approximately this depth, which is likely stronger during summer, when there is more precipitation (Torgersen et al., 1981). Additionally, it is possible that the benthic fluxes of dissolved iron and CO_2_ into the deep monimolimnion are enhanced during the summer through greater sedimentary remineralization due the higher flux of biomass to the lakebed.

### Assemblage partitioning

The chemoorganotrophic aerobes and anaerobes that were common in PA assemblages both metabolize complex organic substrates that are found in particles (e.g., proteins, polysaccharides, chitin, lignin, cellulose), and have particle-adhering abilities. The proportion of taxa that specialize in oxidizing proteinaceous substrates (e.g., Flavobacteriales, Verrucomicrobia, Planctomycetes, DeLong et al., 1993, Duret et al., 2019) and taxa that oxidize or ferment refractory organic carbon substrates (e.g., Kiritimatiellaeota, Burkholderiaceae, van Vliet et al., 2019, Garrity et al., 2015) appear to vary with time-of-year and with redox condition (corresponding to depth). These variables parallel temporal and depth gradients in labile organic carbon availability. The temporal variability of labile organic carbon substrates is likely driven by the seasonality of the photoautotroph blooms (a source of labile organic carbon in summer) versus decaying plant matter (a source of refractory organic carbon in autumn). The decreasing availability of labile organic carbon over depth is likely caused by the continuous remineralization of biogenic debris as they sink towards the lakebed. Our phylum-level analyses revealed that low-abundance chemoorganotrophic environmental candidate phyla belonging to the so-called “microbial dark matter” may collectively also participate in organic carbon remineralization in particles. These phyla have been found attached to particles in marine microbial mats (Armatimonadetes, Burns et al., 2004, Ley et al., 2006) and are enriched in particles in hypoxic and anoxic marine water (Kiritimatiellaeota, Capo et al., 2020). Omics studies have shown that they have surface-adhesion and metabolic abilities similar to better studied taxa that were over-represented in particles in FGL (Eloe-Fadrosh et al., 2016, van Vliet et al., 2019). Other taxa that were over-represented in PA assemblages consume fermentation products like hydrogen and methanol (Hydrogendentes, Methanofastidiosales), and are likely benefiting from the remineralization of organic carbon substrates on particles.

## Conclusion

Oxygen-depleted water bodies like FGL are a window into the geologic past and, likely, the future. As oxygen-deficient waters continue to geographically expand due to global climate change (Stramma et al., 2008, Wright et al., 2012, Moffitt et al., 2015, Breitburg et al., 2018), similar biogeochemical processes are likely to profoundly impact interconnected global biogeochemical cycles (e.g., iron, hydrogen, sulfur, carbon, nitrogen). Ultimately, the remineralization of organic carbon substrates in particles as they sink through distinctive redox zones drives many of these complex interplays. These synergistic processes could potentially contribute to nitrogen deficits through denitrification, enhanced primary production through enriched dissolved iron availability, increased greenhouse gas emissions through methanogenesis, and harm to macrofauna through acidification and hydrogen sulfide production. Therefore, studying the niche-partitioning and drivers of diversity within and between particle-associated and free-living microbial assemblages in these systems is critical.

In permanently redox-stratified FGL, the 16S rRNA amplicon composition of all samples significantly differed among redox conditions and assemblage types, and those from the mixolimnion and oxycline also differed by sampling date. These compositional differences all appear to ultimately be driven by availability and composition of organic carbon substrates and oxidants. Particle-associated organotrophs specialized in surface-adhesion and metabolizing complex organic carbon substrates or fermentation products, whereas free-living chemoorganotrophs seemed to metabolize dissolved amino acids, volatile fatty acids, or hydrocarbons. Variations in organic carbon substrate and oxidant availability above the pycnocline were likely caused by seasonal phenomena, with photoautotrophy being dominant in summer, and wind mixing and potentially allochthonous plant input being more important in autumn. At depth, organic carbon availability may be regulated by vertical transport and remineralization of primary production products from the mixolimnion and light-scattering layer by particle-associated assemblages. The byproducts of remineralized primary production are likely an important source of dissolved organic substrates for the free-living anaerobes in the monimolimnion. Although functionality of taxa in this phylogenetic study was carefully deduced based on thorough literature review and ancillary analyses, we recommend a metagenomic study of FGL PA and FL assemblages to reveal their functional potential through the construction of metagenome assembled genomes (MAGs) as well as targeted biogeochemical experiments for future research efforts.

## Supporting information

supplementary separate files

## Acknowledgements

We thank Dr. Adith Ramamurti, Daniel Garatea, and John Baum for assistance in sample collection during October 2016 and July 2017. We also thank Drs. Robert Aller and Christina Heilbrun for assistance in quantifying hydrogen sulfide. Finally, we thank Mark Wiggins (building manager, School of Marine and Atmospheric Sciences) and Ian Dwyer (PhD candidate, School of Marine and Atmospheric Sciences) for assistance with preparatory work in the machine shop. This research was supported in part by NSF Grant OCE-1259110 and in part by the Wellesley College Fiske Award (to V. Klepac-Ceraj)

## SUPPLEMENTARY MATERIALS

### Treatment of sensor data and chemical metadata

To create a table of metadata values that corresponded to DNA sampling depths, pre-existing chemical data sets and our continuous sensor data required further processing. Pre-cast readings in air and upcasts were removed from continuous sensor data sets and replicated depth readings were averaged. Pre-cast readings were distinguished from the downcast by finding the maximum difference in conductivity within the first two meters of water. The downcast and upcast were distinguished by designating the deepest depth with the earliest time stamp as the true downcast end. Linear splines were then fit to data points in pre-existing chemical data sets and cleaned, averaged continuous sensor data sets. Continuous sensor data splines were used to solve for the value of the variable at each DNA sampling depth and the chemical data splines were used to solve for values at each DNA sampling depth if a discrete sample was not taken at that exact depth.

### Tables

**Table S1.** Kruskal-Wallace pairwise tests of Pielou-evenness alpha diversity against metadata bins. PS=Particulate sulfur; Phyco=phycoerythrin; ORP= oxidation reduction potential; ox. cat=oxygen/redox category; ICA=inorganic carbon assimilation; BHP=bacterial heterotrophic production; Temp=temperature. If no units are specified, biological and chemical data is concentration.

| *variable* | *Group 1* | *Group 2* | *p* | *q* |
| --- | --- | --- | --- | --- |
| *0.2-0.4 μm PS* | < 0.045 (n=8) | >= 0.045 (n=8) | 0.67 | 0.67 |
| *> 0.4 μm PS* | < 0.01 (n=8) | >= 0.01 < 0.05 (n=6) | 0.70 | 1.00 |
|  | < 0.01 (n=8) | >= 0.05 (n=2) | 1.00 | 1.00 |
|  | >= 0.01 < 0.05 (n=6) | >= 0.05 (n=2) | 0.74 | 1.00 |
| *BHP (μmoles C/L/d)* | < 0.4 (n=16) | >= 0.4 < 1.1 (n=10) | 0.49 | 0.49 |
|  | < 0.4 (n=16) | >= 1.1 (n=4) | 0.07 | 0.11 |
|  | >= 0.4 < 1.1 (n=10) | >= 1.1 (n=4) | 0.05 | 0.11 |
| *CH<sub>4</sub>* | < 2.4 (n=18) | >= 13 (n=4) | 0.55 | 0.66 |
|  | < 2.4 (n=18) | >= 2.4 < 7.5 (n=8) | 0.07 | 0.20 |
|  | < 2.4 (n=18) | >= 7.5 < 13 (n=6) | 0.32 | 0.48 |
|  | >= 2.4 < 7.5 (n=8) | >= 13 (n=4) | 0.13 | 0.25 |
|  | >= 2.4 < 7.5 (n=8) | >= 7.5 < 13 (n=6) | 0.03 | 0.17 |
|  | >= 7.5 < 13 (n=6) | >= 13 (n=4) | 0.67 | 0.67 |
| *Chl-a* | < 3.4 (n=30) | >= 3.4 < 7.4 (n=4) | 0.03 | 0.10 |
|  | < 3.4 (n=30) | >= 7.4 (n=2) | 0.10 | 0.15 |
|  | >= 3.4 < 7.4 (n=4) | >= 7.4 (n=2) | 1.00 | 1.00 |
| $\delta^{13}C_{POC}$ | < -41 (n=2) | >= -35 (n=3) | 0.08 | 0.17 |
|  | < -41 (n=2) | >= -38.5 < -35 (n=8) | 0.19 | 0.23 |
|  | < -41 (n=2) | >= -41 < -38.5 (n=6) | 0.05 | 0.17 |
|  | >= -38.5 < -35 (n=8) | >= -35 (n=3) | 0.07 | 0.17 |
|  | >= -41 < -38.5 (n=6) | >= -35 (n=3) | 0.12 | 0.18 |
|  | >= -41 < -38.5 (n=6) | >= -38.5 < -35 (n=8) | 0.25 | 0.25 |
| *Depth (m)* | < 14 (n=4) | >= 14 < 30 (n=28) | 0.25 | 0.38 |
|  | < 14 (n=4) | >= 30 (n=4) | 0.15 | 0.38 |
|  | >= 14 < 30 (n=28) | >= 30 (n=4) | 0.61 | 0.61 |
| *Fe* | < 216 (n=24) | >= 1000 (n=2) | 0.44 | 0.63 |
|  | < 216 (n=24) | >= 216 < 425 (n=6) | 0.64 | 0.80 |
|  | < 216 (n=24) | >= 425 < 600 (n=2) | 0.34 | 0.63 |
|  | < 216 (n=24) | >= 600 < 1000 (n=2) | 0.15 | 0.50 |
|  | >= 216 < 425 (n=6) | >= 1000 (n=2) | 0.32 | 0.63 |
|  | >= 216 < 425 (n=6) | >= 425 < 600 (n=2) | 0.74 | 0.82 |
|  | >= 216 < 425 (n=6) | >= 600 < 1000 (n=2) | 0.05 | 0.46 |
|  | >= 425 < 600 (n=2) | >= 1000 (n=2) | 0.44 | 0.63 |
|  | >= 425 < 600 (n=2) | >= 600 < 1000 (n=2) | 0.12 | 0.50 |
|  | >= 600 < 1000 (n=2) | >= 1000 (n=2) | 1.00 | 1.00 |
| *ICA (μmoles*<br>*C/L/d)* | < 1 (n=16) | >= 1 < 8 (n=10) | 0.92 | 0.92 |
|  | < 1 (n=16) | >= 8 (n=4) | 0.05 | 0.14 |
|  | >= 1 < 8 (n=10) | >= 8 (n=4) | 0.26 | 0.39 |
| *Mn* | < 8000 (n=18) | >= 30000 < 52000 (n=4) | 0.27 | 0.53 |
|  | < 8000 (n=18) | >= 52000 (n=2) | 0.10 | 0.43 |
|  | < 8000 (n=18) | >= 8000 < 30000 (n=12) | 0.97 | 0.97 |
|  | >= 30000 < 52000 (n=4) | >= 52000 (n=2) | 0.35 | 0.53 |
|  | >= 8000 < 30000 (n=12) | >= 30000 < 52000 (n=4) | 0.63 | 0.75 |
|  | >= 8000 < 30000 (n=12) | >= 52000 (n=2) | 0.14 | 0.43 |
| *Month* | early October (n=16) | late July (n=20) | 0.24 | 0.24 |
| *NH<sub>4</sub>* | < 50 (n=6) | >= 150 < 325 (n=4) | 0.39 | 0.59 |
|  | < 50 (n=6) | >= 325 (n=2) | 0.50 | 0.60 |
|  | < 50 (n=6) | >= 50 < 150 (n=8) | 0.05 | 0.19 |
|  | >= 150 < 325 (n=4) | >= 325 (n=2) | 0.16 | 0.33 |
|  | >= 50 < 150 (n=8) | >= 150 < 325 (n=4) | 0.06 | 0.19 |
|  | >= 50 < 150 (n=8) | >= 325 (n=2) | 0.60 | 0.60 |
| *NO<sub>3</sub><sup>-</sup>* | < 6 (n=18) | >= 25 < 50 (n=8) | 0.29 | 0.87 |
|  | < 6 (n=18) | >= 50 (n=4) | 0.50 | 0.99 |
|  | < 6 (n=18) | >= 6 < 25 (n=6) | 1.00 | 1.00 |
|  | >= 25 < 50 (n=8) | >= 50 (n=4) | 0.13 | 0.76 |
|  | >= 6 < 25 (n=6) | >= 25 < 50 (n=8) | 0.80 | 1.00 |
|  | >= 6 < 25 (n=6) | >= 50 (n=4) | 0.83 | 1.00 |
| *O<sub>2</sub>* | < 3.5 (n=10) | >= 250 < 500 (n=6) | 0.91 | 1.00 |
|  | < 3.5 (n=10) | >= 3.5 < 22 (n=14) | 0.95 | 1.00 |
|  | < 3.5 (n=10) | >= 500 (n=2) | 0.52 | 1.00 |
|  | < 3.5 (n=10) | >= 90 < 250 (n=4) | 0.57 | 1.00 |
|  | >= 250 < 500 (n=6) | >= 500 (n=2) | 0.74 | 1.00 |
|  | >= 3.5 < 22 (n=14) | >= 250 < 500 (n=6) | 0.80 | 1.00 |
|  | >= 3.5 < 22 (n=14) | >= 500 (n=2) | 0.75 | 1.00 |
|  | >= 3.5 < 22 (n=14) | >= 90 < 250 (n=4) | 0.60 | 1.00 |
|  | >= 90 < 250 (n=4) | >= 250 < 500 (n=6) | 0.39 | 1.00 |
|  | >= 90 < 250 (n=4) | >= 500 (n=2) | 1.00 | 1.00 |
| *ORP (mV)* | < -350 (n=12) | >= -180 < 0 (n=2) | 0.20 | 0.71 |
|  | < -350 (n=12) | >= -350 < -180 (n=4) | 1.00 | 1.00 |
|  | < -350 (n=12) | >= 0 (n=18) | 0.97 | 1.00 |
|  | >= -180 < 0 (n=2) | >= 0 (n=18) | 0.31 | 0.71 |
|  | >= -350 < -180 (n=4) | >= -180 < 0 (n=2) | 0.35 | 0.71 |
|  | >= -350 < -180 (n=4) | >= 0 (n=18) | 1.00 | 1.00 |
| *Oxycline pos.* | above (n=4) | below (n=14) | 0.29 | 0.54 |
|  | above (n=4) | inside (n=10) | 0.32 | 0.54 |
|  | above (n=4) | lower boundary (n=4) | 0.77 | 0.86 |
|  | above (n=4) | upper boundary (n=4) | 0.04 | 0.22 |
|  | below (n=14) | inside (n=10) | 0.95 | 0.95 |
|  | below (n=14) | lower boundary (n=4) | 0.29 | 0.54 |
|  | below (n=14) | upper boundary (n=4) | 0.52 | 0.71 |
|  | inside (n=10) | lower boundary (n=4) | 0.26 | 0.54 |
|  | inside (n=10) | upper boundary (n=4) | 0.57 | 0.71 |
|  | lower boundary (n=4) | upper boundary (n=4) | 0.04 | 0.22 |
| *pH* | < 6.87 (n=14) | >= 6.87 < 7.16 (n=12) | 0.72 | 0.97 |
|  | < 6.87 (n=14) | >= 7.16 < 7.47 (n=6) | 0.87 | 0.97 |
|  | < 6.87 (n=14) | >= 7.47 < 7.73 (n=2) | 0.20 | 0.51 |
|  | < 6.87 (n=14) | >= 7.73 (n=2) | 0.75 | 0.97 |
|  | >= 6.87 < 7.16 (n=12) | >= 7.16 < 7.47 (n=6) | 1.00 | 1.00 |
|  | >= 6.87 < 7.16 (n=12) | >= 7.47 < 7.73 (n=2) | 0.14 | 0.48 |
|  | >= 6.87 < 7.16 (n=12) | >= 7.73 (n=2) | 0.86 | 0.97 |
|  | >= 7.16 < 7.47 (n=6) | >= 7.47 < 7.73 (n=2) | 0.05 | 0.46 |
|  | >= 7.16 < 7.47 (n=6) | >= 7.73 (n=2) | 0.74 | 0.97 |
|  | >= 7.47 < 7.73 (n=2) | >= 7.73 (n=2) | 0.12 | 0.48 |
| *Phyco.* | < 13.5 (n=28) | >= 128 (n=2) | 0.13 | 0.32 |
|  | < 13.5 (n=28) | >= 13.5 < 128 (n=6) | 0.28 | 0.32 |
|  | >= 13.5 < 128 (n=6) | >= 128 (n=2) | 0.32 | 0.32 |
| *PS* | < 1 (n=30) | >= 1 < 11 (n=4) | 0.08 | 0.12 |
|  | < 1 (n=30) | >= 11 (n=2) | 0.07 | 0.12 |
|  | >= 1 < 11 (n=4) | >= 11 (n=2) | 1.00 | 1.00 |
| *Redox* | euxinic (n=18) | hypoxic (n=2) | 0.31 | 0.60 |
|  | euxinic (n=18) | oxic (n=12) | 0.64 | 0.64 |
|  | euxinic (n=18) | suboxic (n=4) | 0.50 | 0.60 |
|  | hypoxic (n=2) | oxic (n=12) | 0.36 | 0.60 |
|  | hypoxic (n=2) | suboxic (n=4) | 0.35 | 0.60 |
|  | oxic (n=12) | suboxic (n=4) | 0.47 | 0.60 |
| *Salinity (psu)* | < 0.9 (n=2) | >= 0.9 < 1.22 (n=12) | 0.07 | 0.18 |
|  | < 0.9 (n=2) | >= 1.22 (n=22) | 0.12 | 0.18 |
|  | >= 0.9 < 1.22 (n=12) | >= 1.22 (n=22) | 0.43 | 0.43 |
| *SizeFrac* | 0.2-2.7 (n=18) | >2.7 (n=18) | 0.45 | 0.45 |
| *SO<sub>4</sub><sup>2-</sup>* | < 13 (n=26) | >= 13 (n=10) | 0.09 | 0.09 |
| *Sulfide* | < 400 (n=20) | >= 2000 (n=2) | 0.30 | 0.51 |
|  | < 400 (n=20) | >= 400 < 2000 (n=14) | 0.97 | 0.97 |
|  | >= 400 < 2000 (n=14) | >= 2000 (n=2) | 0.34 | 0.51 |
| *TDS* | < 1550 (n=12) | >= 1550 < 1766.7 (n=20) | 0.51 | 0.54 |
|  | < 1550 (n=12) | >= 1766.7 (n=4) | 0.54 | 0.54 |
|  | >= 1550 < 1766.7 (n=20) | >= 1766.7 (n=4) | 0.49 | 0.54 |
| *Temp (°C)* | < 7.2 (n=2) | >= 15 (n=2) | 0.12 | 0.23 |
|  | < 7.2 (n=2) | >= 7.2 < 8.3 (n=8) | 0.19 | 0.23 |
|  | < 7.2 (n=2) | >= 8.3 < 15 (n=24) | 0.08 | 0.23 |
|  | >= 7.2 < 8.3 (n=8) | >= 15 (n=2) | 0.04 | 0.22 |
|  | >= 7.2 < 8.3 (n=8) | >= 8.3 < 15 (n=24) | 0.38 | 0.38 |
|  | >= 8.3 < 15 (n=24) | >= 15 (n=2) | 0.18 | 0.23 |
| *Thiosulfate* | < 0.5 (n=26) | >= 0.5 (n=10) | 0.92 | 0.92 |
| *Total cells* | < 2.0e+09 (n=2) | >= 1.2e+10 (n=2) | 1.00 | 1.00 |
|  | < 2.0e+09 (n=2) | >= 2.0e+09 < 7.0e+09 (n=22) | 0.06 | 0.18 |
|  | < 2.0e+09 (n=2) | >= 7.0e+09 < 1.2e+10 (n=4) | 0.35 | 0.53 |
|  | >= 2.0e+09 < 7.0e+09 (n=22) | >= 1.2e+10 (n=2) | 0.06 | 0.18 |
|  | >= 2.0e+09 < 7.0e+09 (n=22) | >= 7.0e+09 < 1.2e+10 (n=4) | 0.32 | 0.53 |
|  | >= 7.0e+09 < 1.2e+10 (n=4) | >= 1.2e+10 (n=2) | 0.64 | 0.77 |
| *Turbidity (FNU)* | < 0.5 (n=18) | >= 0.5 < 1.35 (n=4) | 0.44 | 0.53 |
|  | < 0.5 (n=18) | >= 1.35 < 35 (n=12) | 0.29 | 0.53 |
|  | < 0.5 (n=18) | >= 35 (n=2) | 0.26 | 0.53 |
|  | >= 0.5 < 1.35 (n=4) | >= 1.35 < 35 (n=12) | 0.90 | 0.90 |
|  | >= 0.5 < 1.35 (n=4) | >= 35 (n=2) | 0.35 | 0.53 |
|  | >= 1.35 < 35 (n=12) | >= 35 (n=2) | 0.27 | 0.53 |
| *Year* | 2016.0 (n=16) | 2017.0 (n=20) | 0.24 | 0.24 |

**Table S2.** Permutational analysis of variance (PERMANOVA) performed on the Bray-Curtis dissimilarity. PS=Particulate sulfur; Phyco=phycoerythrin; ORP= oxidation reduction potential; ox. cat=oxygen/redox category; ICA=inorganic carbon assimilation; BHP=bacterial heterotrophic production; TDS=total dissolved solids; Temp=temperature. If no units are specified, biological and chemical data is concentration.

| *variable* | *sample*<br>*size* | *no.*<br>*groups* | *p* |
| --- | --- | --- | --- |
| *Fe* | 36 | 5 | 0.001 |
| *CH<sub>4</sub>* | 36 | 4 | 0.001 |
| *NH<sub>4</sub>* | 20 | 4 | 0.001 |
| *BHP (μmoles C/L/d)* | 30 | 3 | 0.001 |
| *Thiosulfate* | 36 | 2 | 0.001 |
| *TDS* | 36 | 3 | 0.001 |
| *Salinity (psu)* | 36 | 3 | 0.001 |
| *Mn* | 36 | 4 | 0.001 |
| *Oxcat* | 36 | 4 | 0.001 |
| *pH* | 36 | 5 | 0.001 |
| *ORP (mV)* | 36 | 4 | 0.001 |
| *Temp (°C)* | 36 | 4 | 0.001 |
| *Turbidity (FNU)* | 36 | 4 | 0.001 |
| *O<sub>2</sub>* | 36 | 5 | 0.001 |
| *Sulfide* | 36 | 3 | 0.001 |
| *oxyclinepos* | 36 | 5 | 0.001 |
| *SO<sub>4</sub><sup>2-</sup>* | 36 | 2 | 0.001 |
| *NO<sub>3</sub><sup>-</sup>* | 36 | 4 | 0.001 |
| *Phyco.* | 36 | 3 | 0.002 |
| *Depth (m)* | 36 | 3 | 0.003 |
| *Total cells* | 30 | 4 | 0.005 |
| *ICA (μmoles C/L/d)* | 30 | 3 | 0.005 |
| *PS* | 36 | 3 | 0.007 |
| *SizeFrac* | 36 | 2 | 0.013 |
| *δ<sup>13</sup>C<sub>POC</sub>* | 19 | 4 | 0.019 |
| *Month* | 36 | 2 | 0.03 |
| *Year* | 36 | 2 | 0.033 |
| *> 0.4 μm PS* | 16 | 3 | 0.079 |
| *Chl-a* | 36 | 3 | 0.147 |
| *0.2-0.4 μm PS* | 16 | 2 | 0.18 |

### Figures

In all figures below, broken lines represent oxycline boundaries. Figures A.1-A.9 show measurements acquired with a YSI EXO1 sonde.

**Figure S1.**
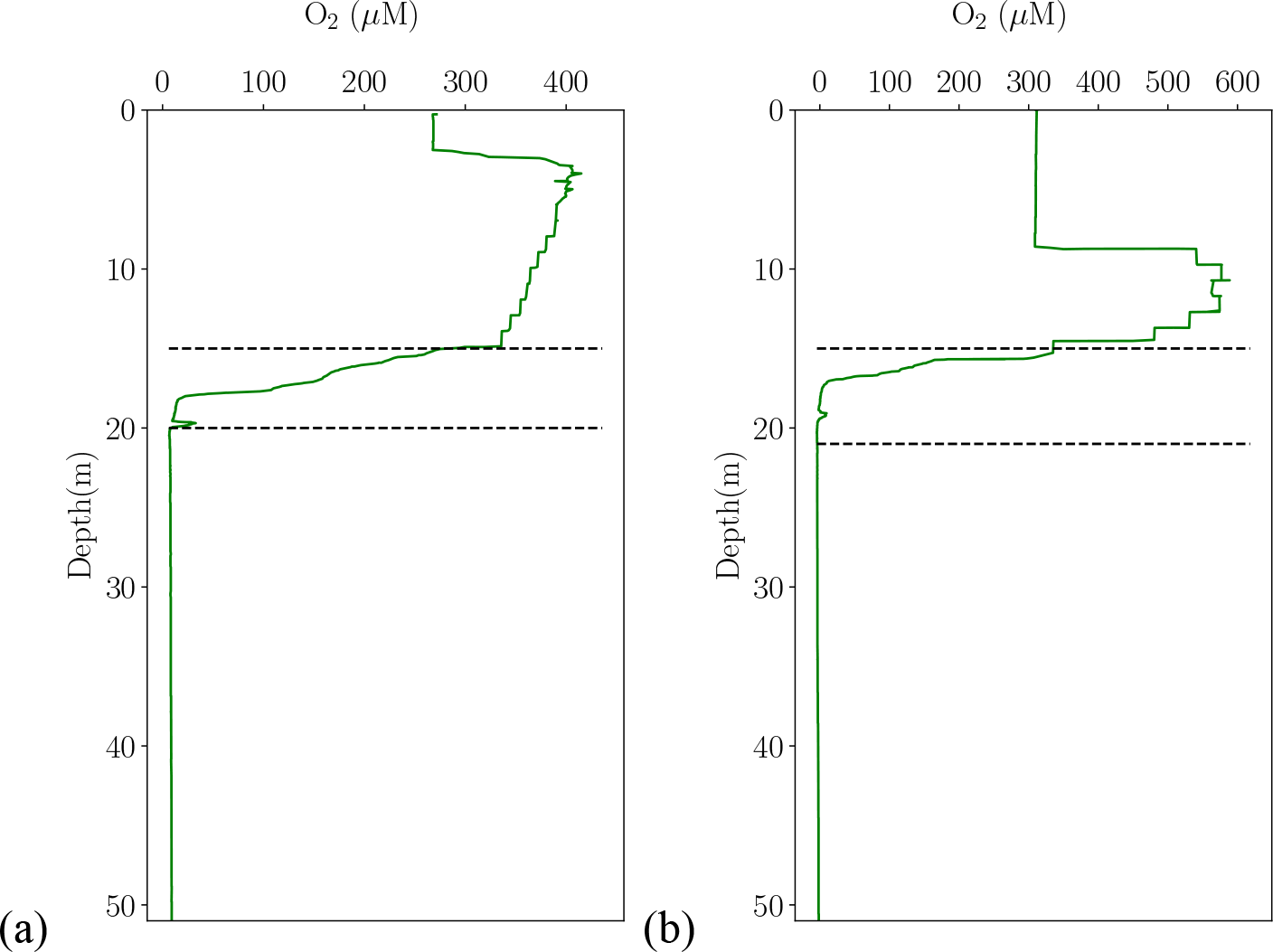
Oxygen concentration in (a) July and (b) October.

**Figure S2.**
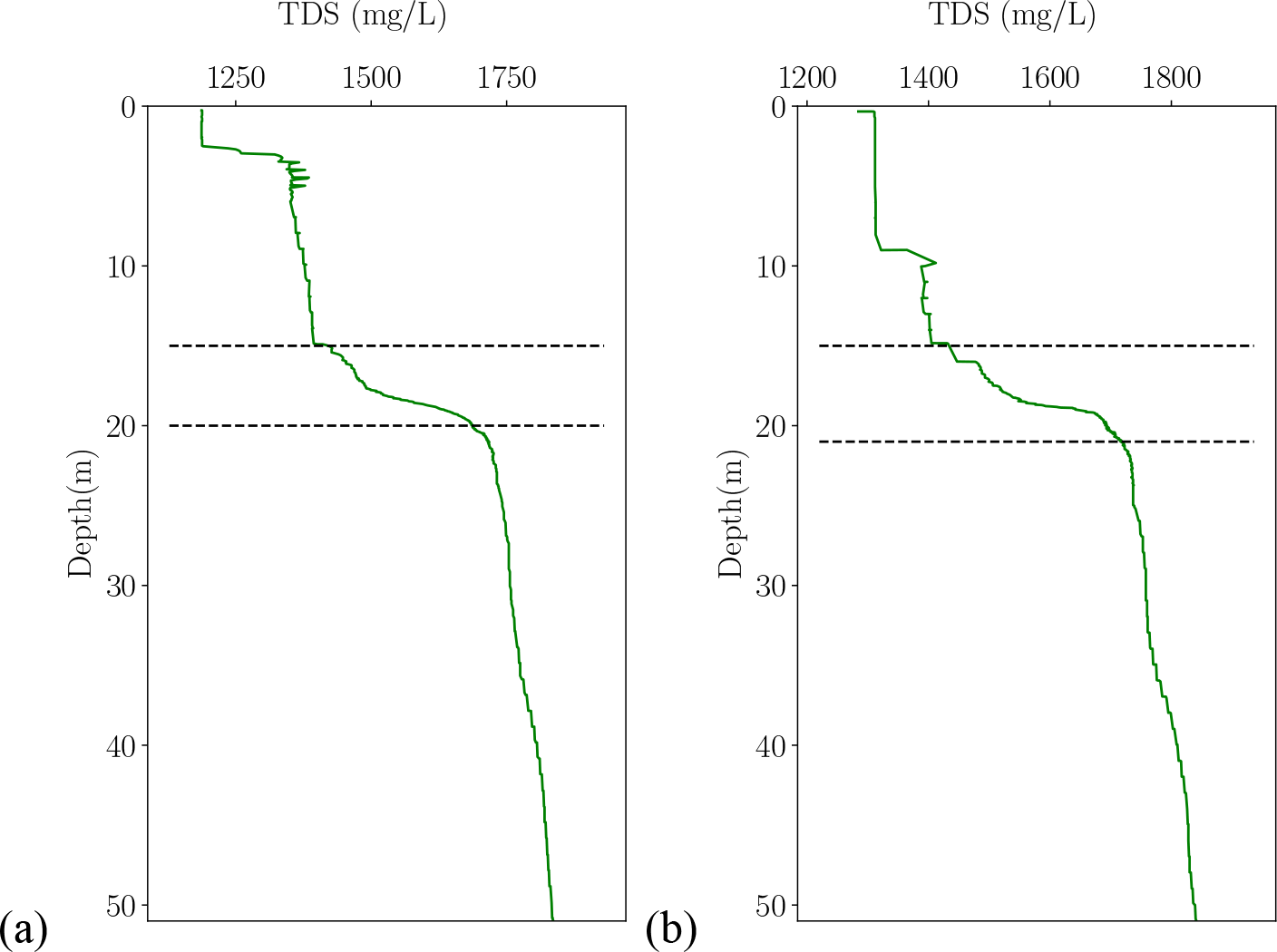
Total Dissolved Solids (TDS) in (a) July and (b) October.

**Figure S3.**
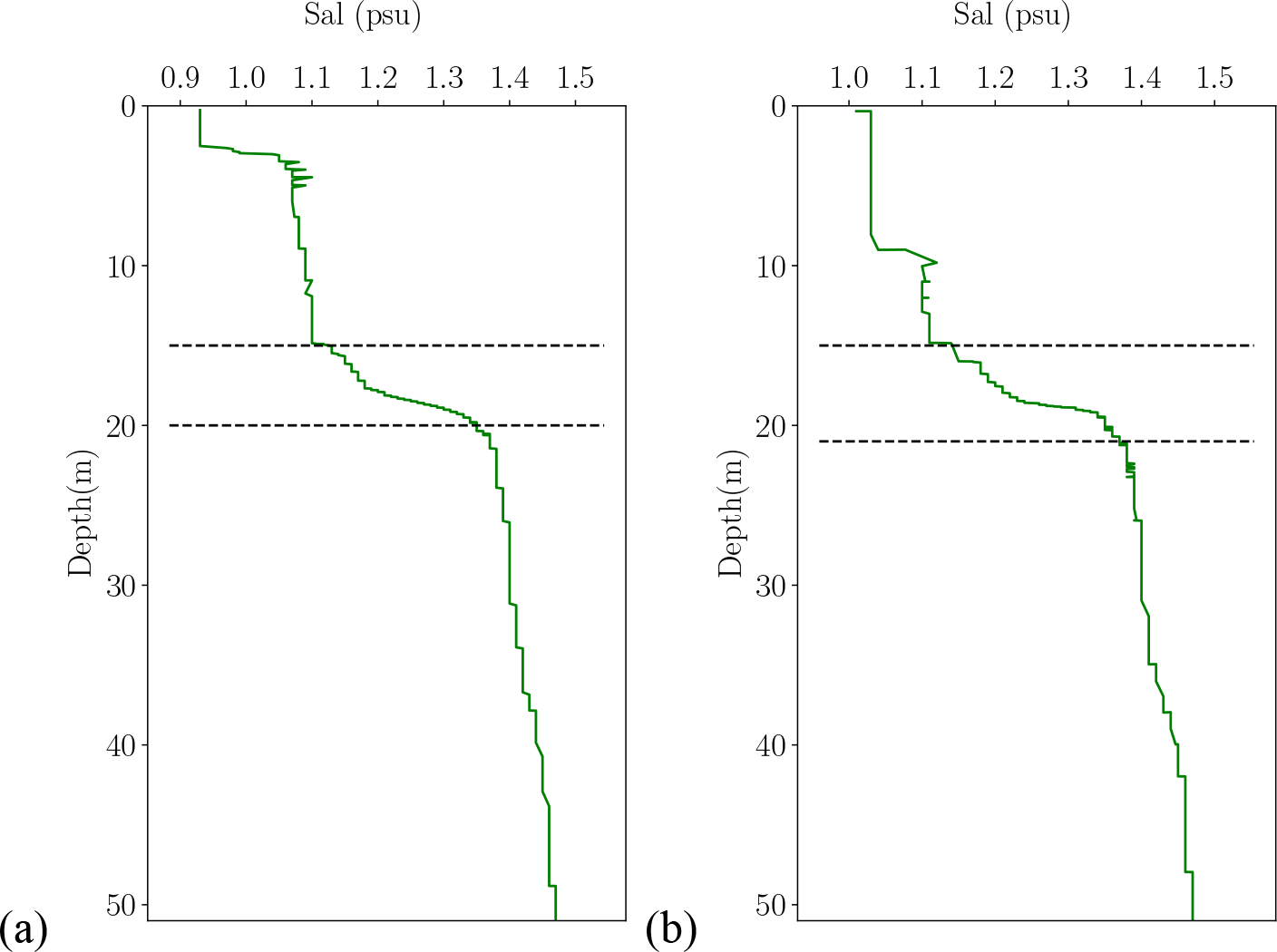
Salinity in (a) July and (b) October.

**Figure S4.**
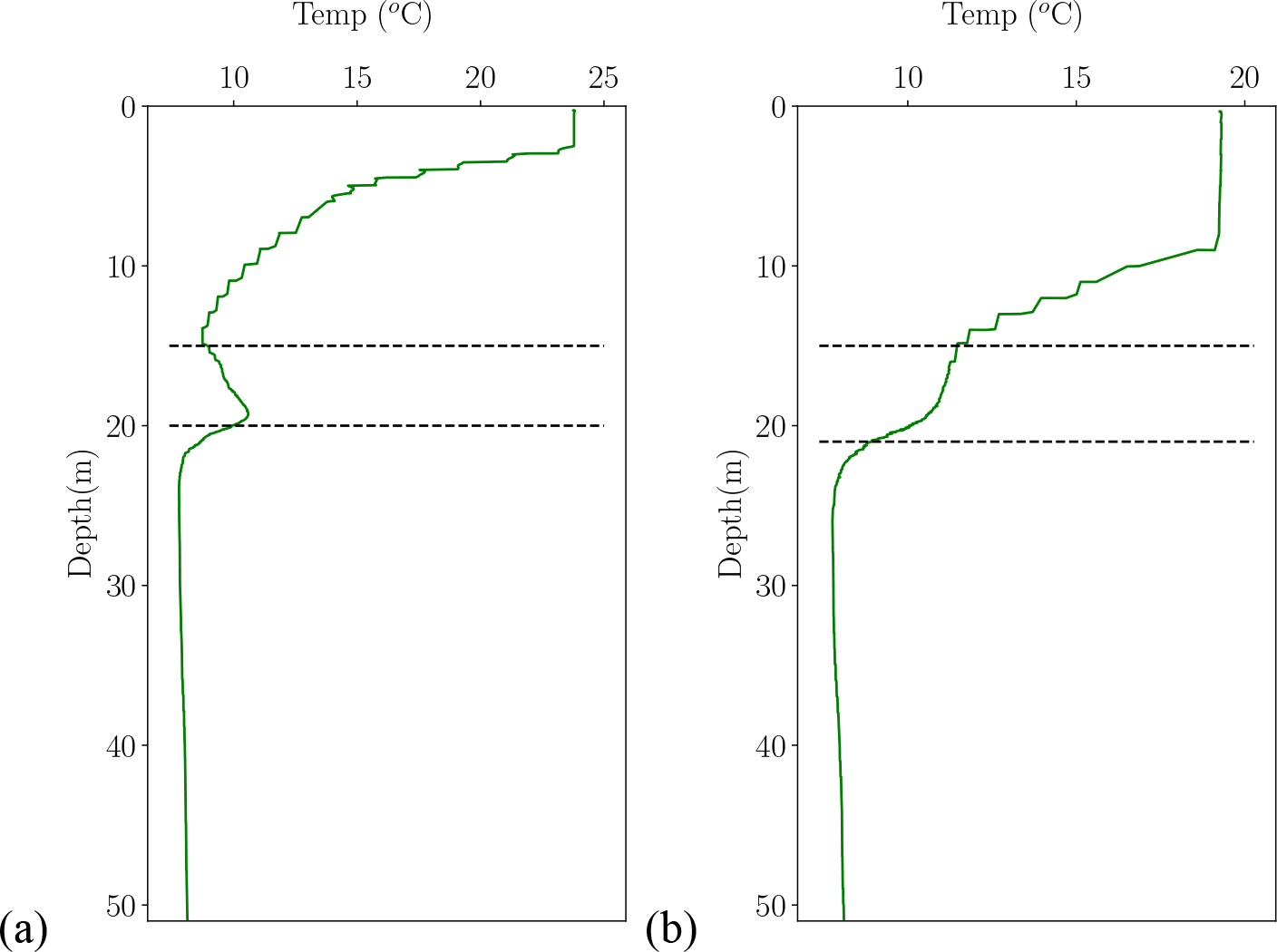
Temperature in (a) July and (b) October.

**Figure S5.**
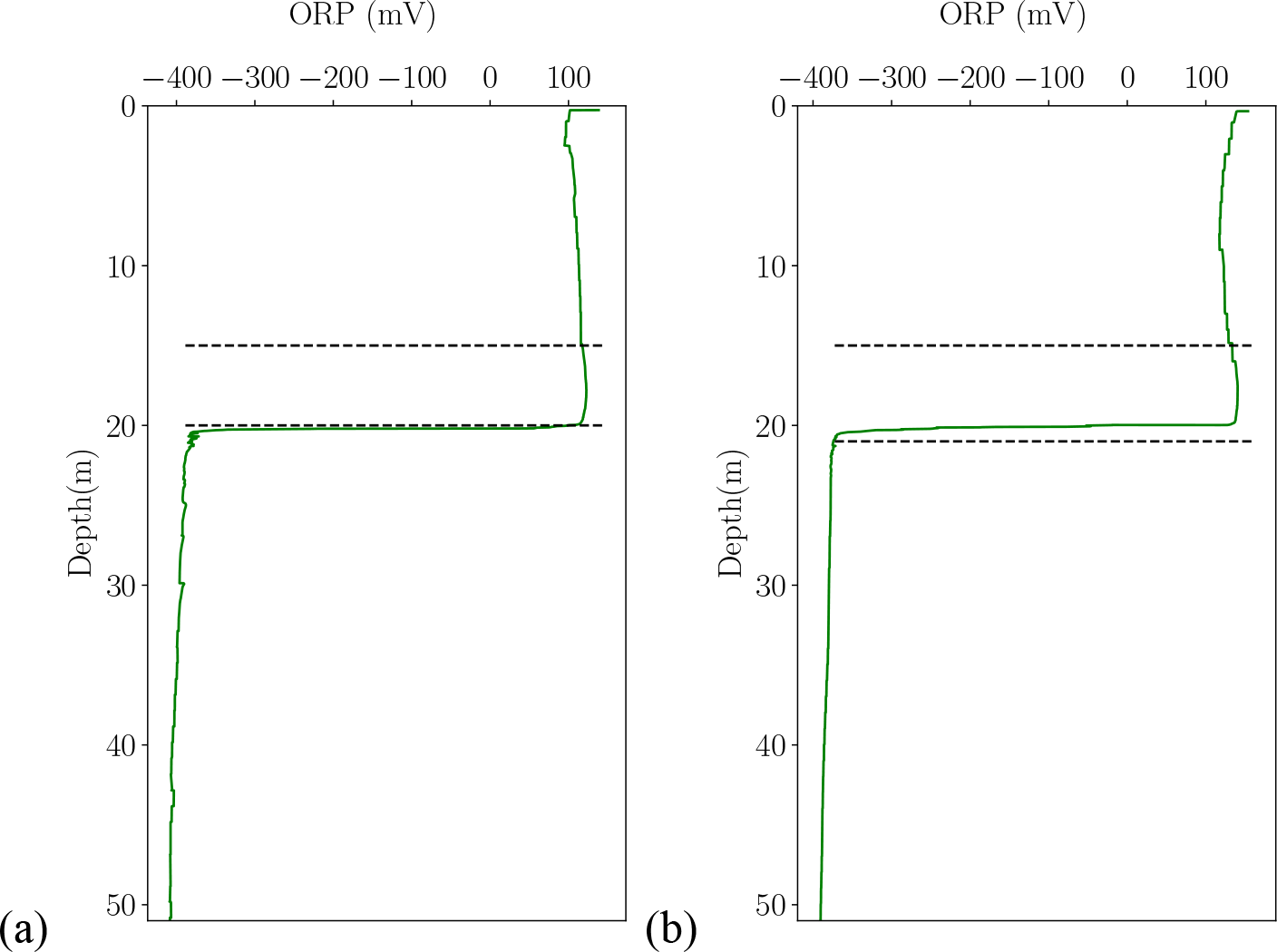
Oxidation reduction potential (ORP) in (a) July and (b) October.

**Figure S6.**
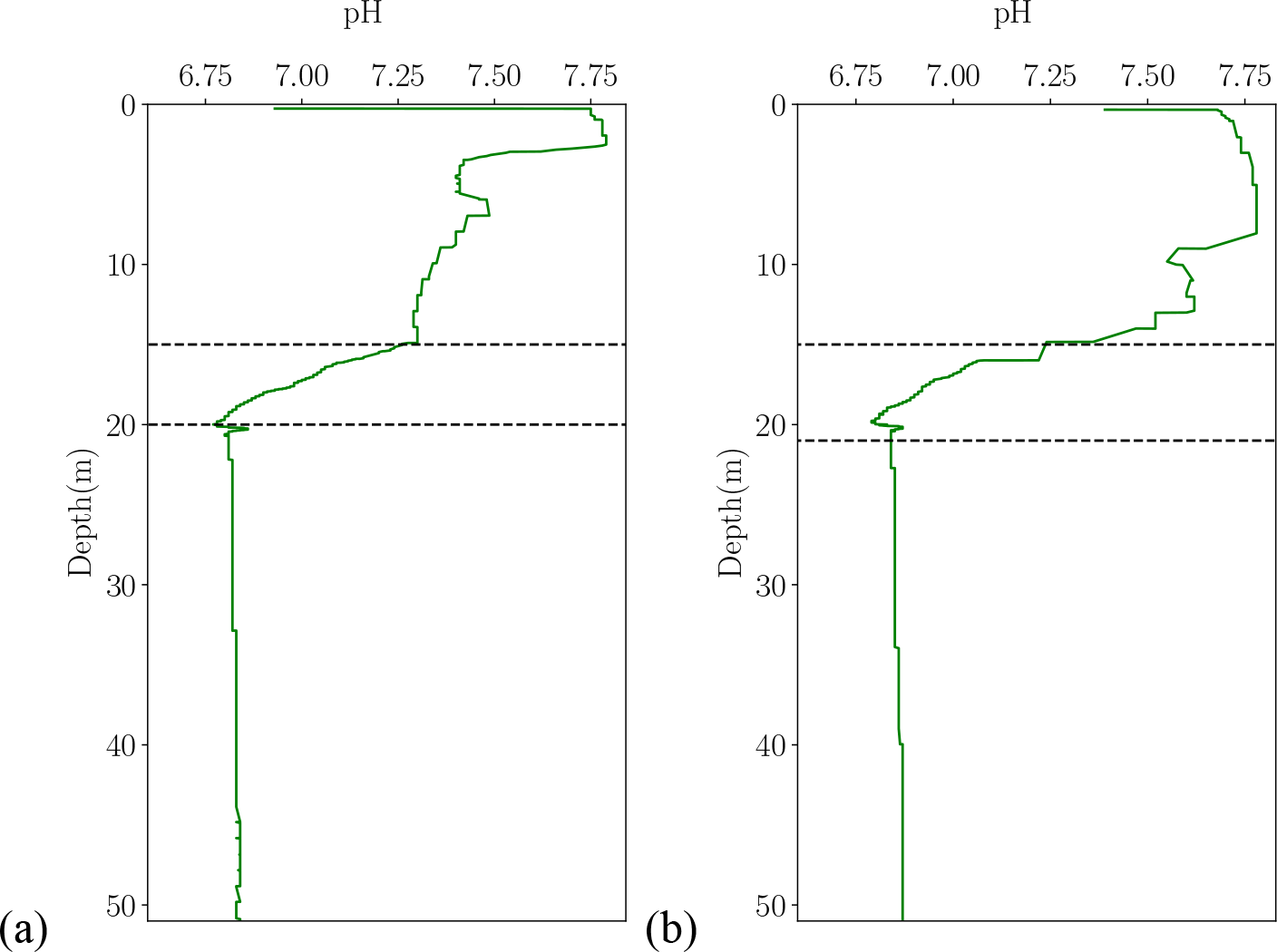
pH in (a) July and (b) October.

**Figure S7.**
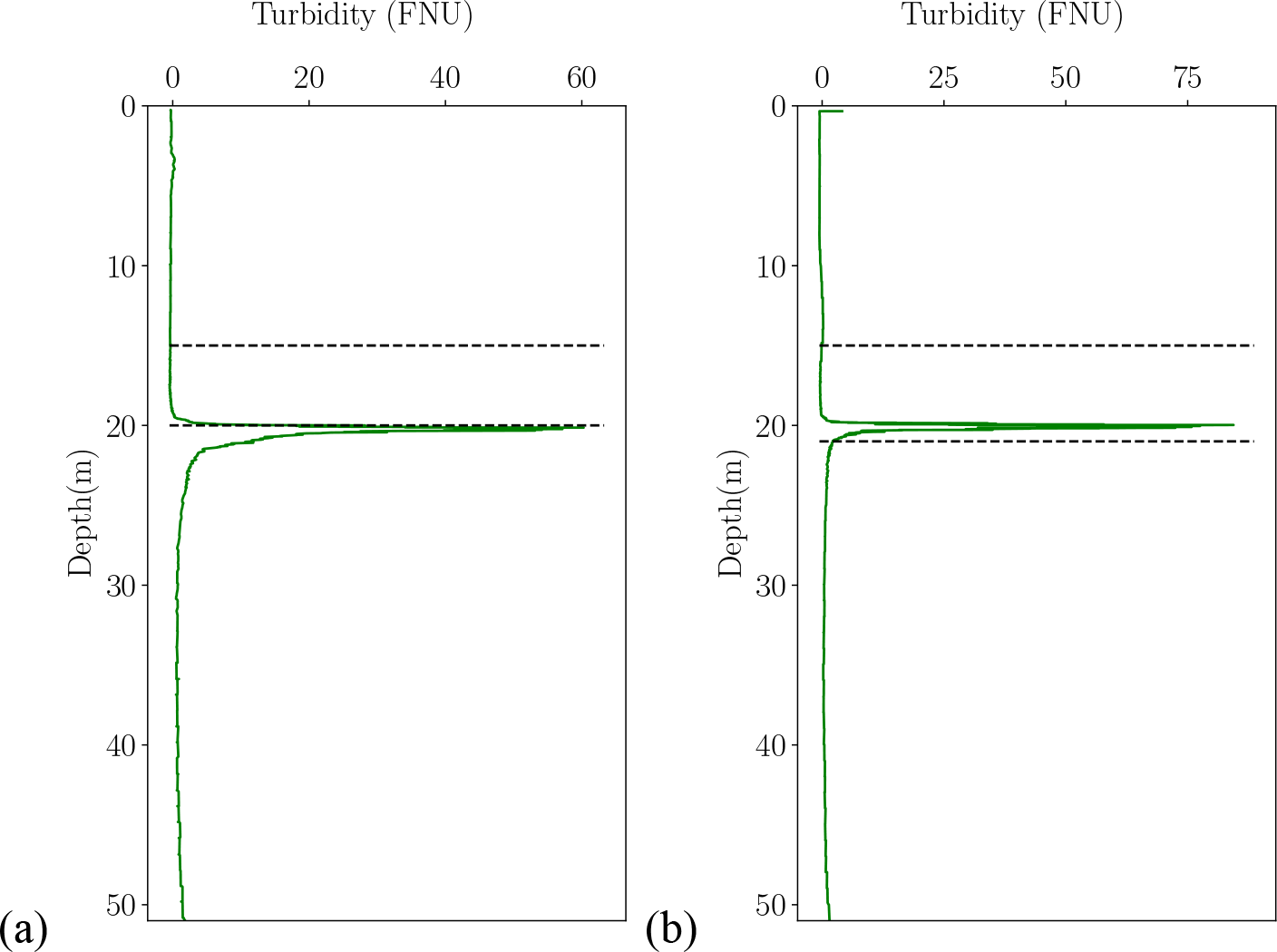
Turbidity in (a) July and (b) October.

**Figure S8.**
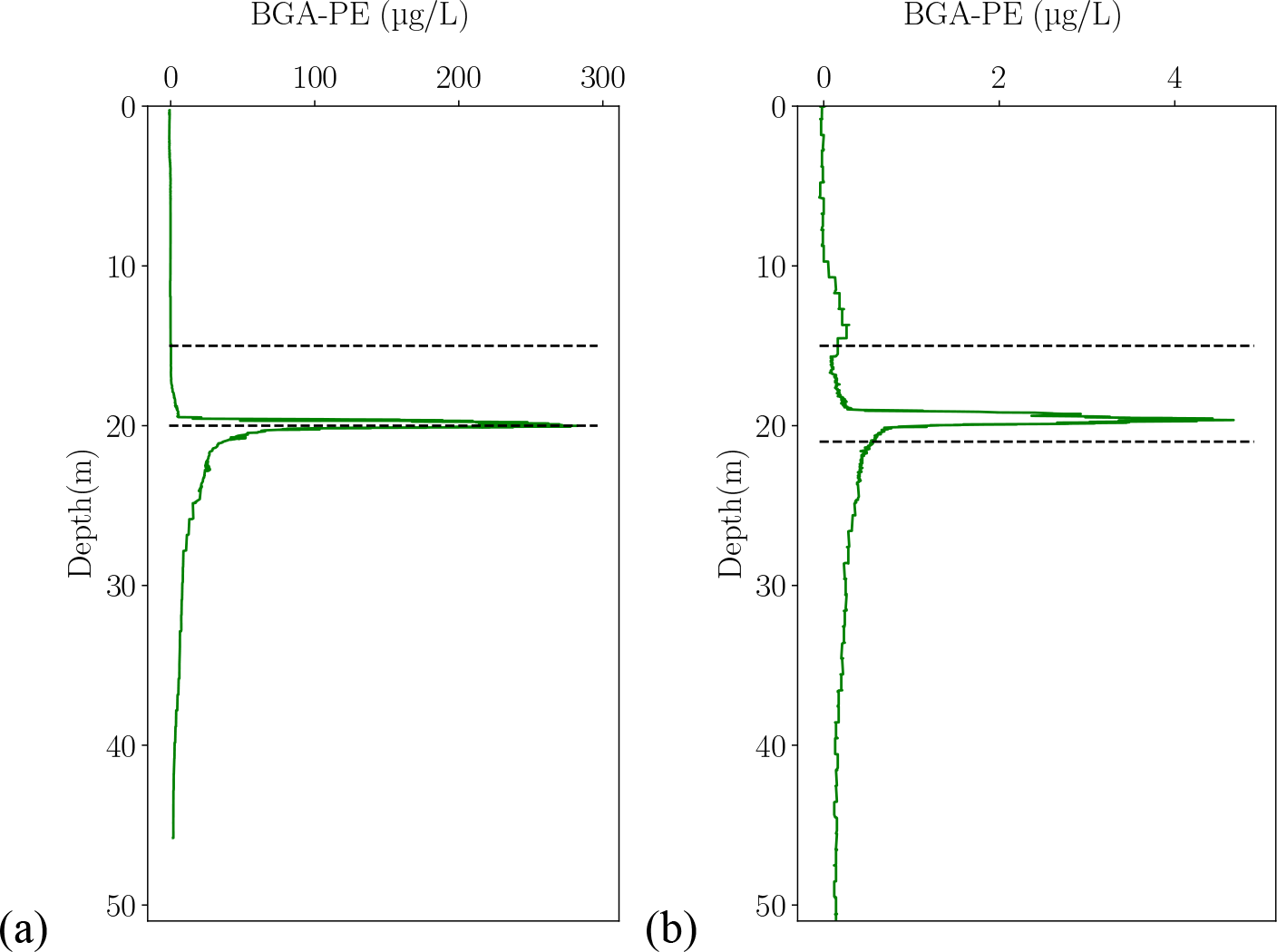
Phycoerythrin concentration in (a) July and (b) October.

**Figure S9.**
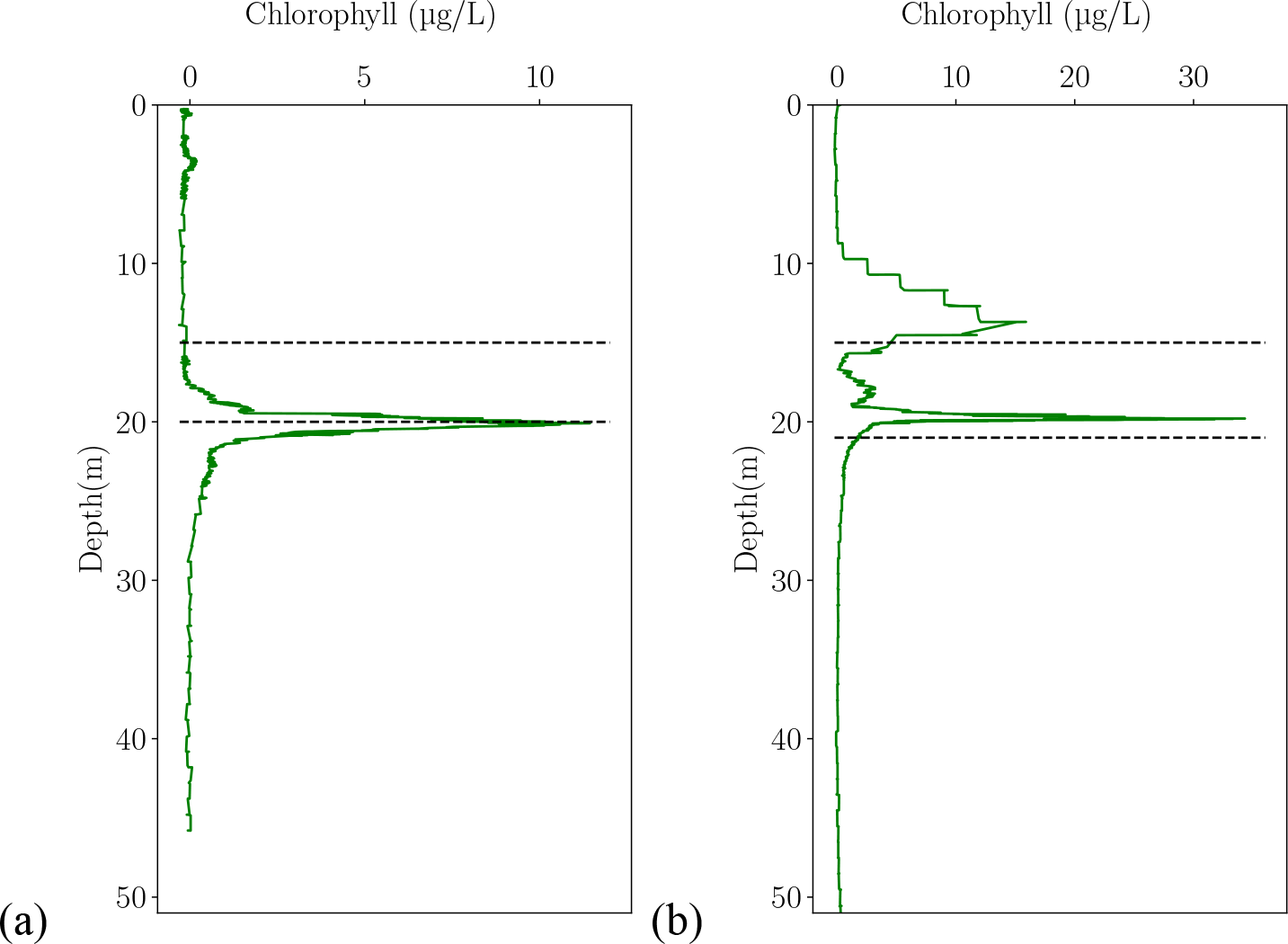
Chlorophyll-a concentration in (a) July and (b) October.

**Figure S10.**
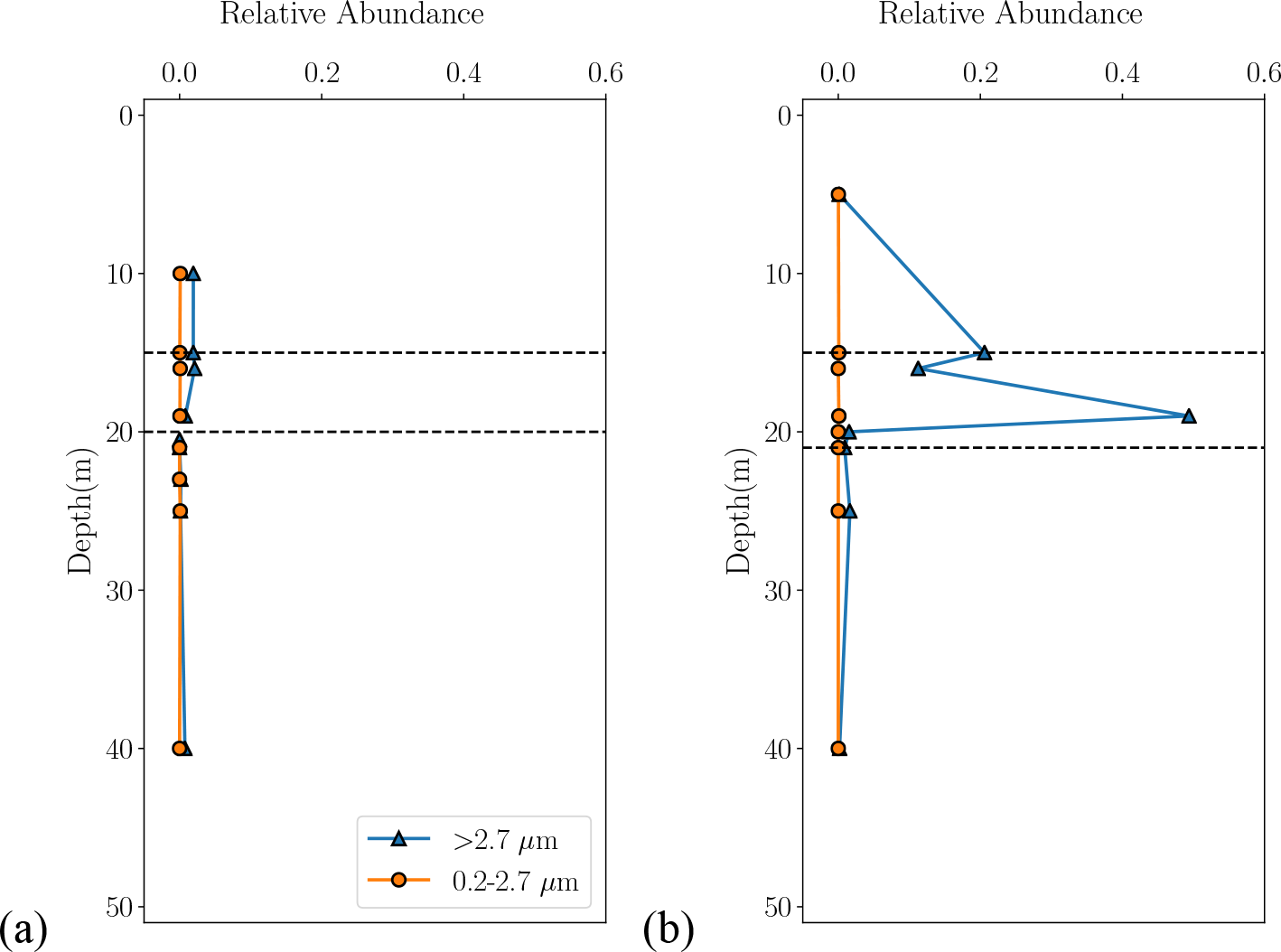
The relative abundance of all chlorophyte mitochondrion and chloroplast 16S DNA sequences present in the >2.7 μm (PA) and 0.2-2.7 μm (FL) size fractions in 16S rRNA libraries, for (a) July and (b) October.

