## Supplementary figures and images for "Particle-associated and free-living microbial assemblages are distinct in a permanently redox-stratified freshwater lake"

### CohenA_ORfactor_categorical_genuslevel.pdf

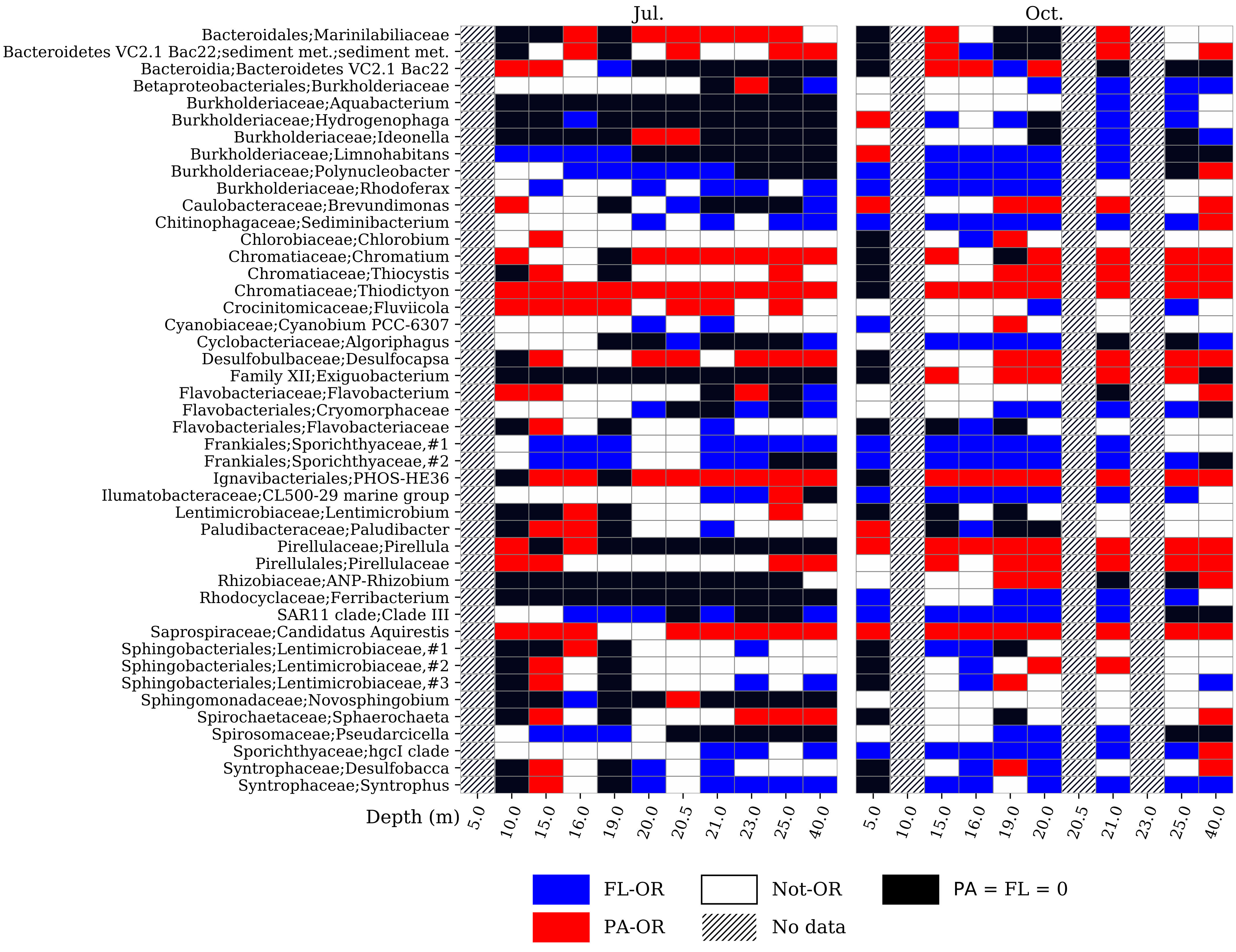

### CohenA_ORfactor_categorical_orderlevel.pdf

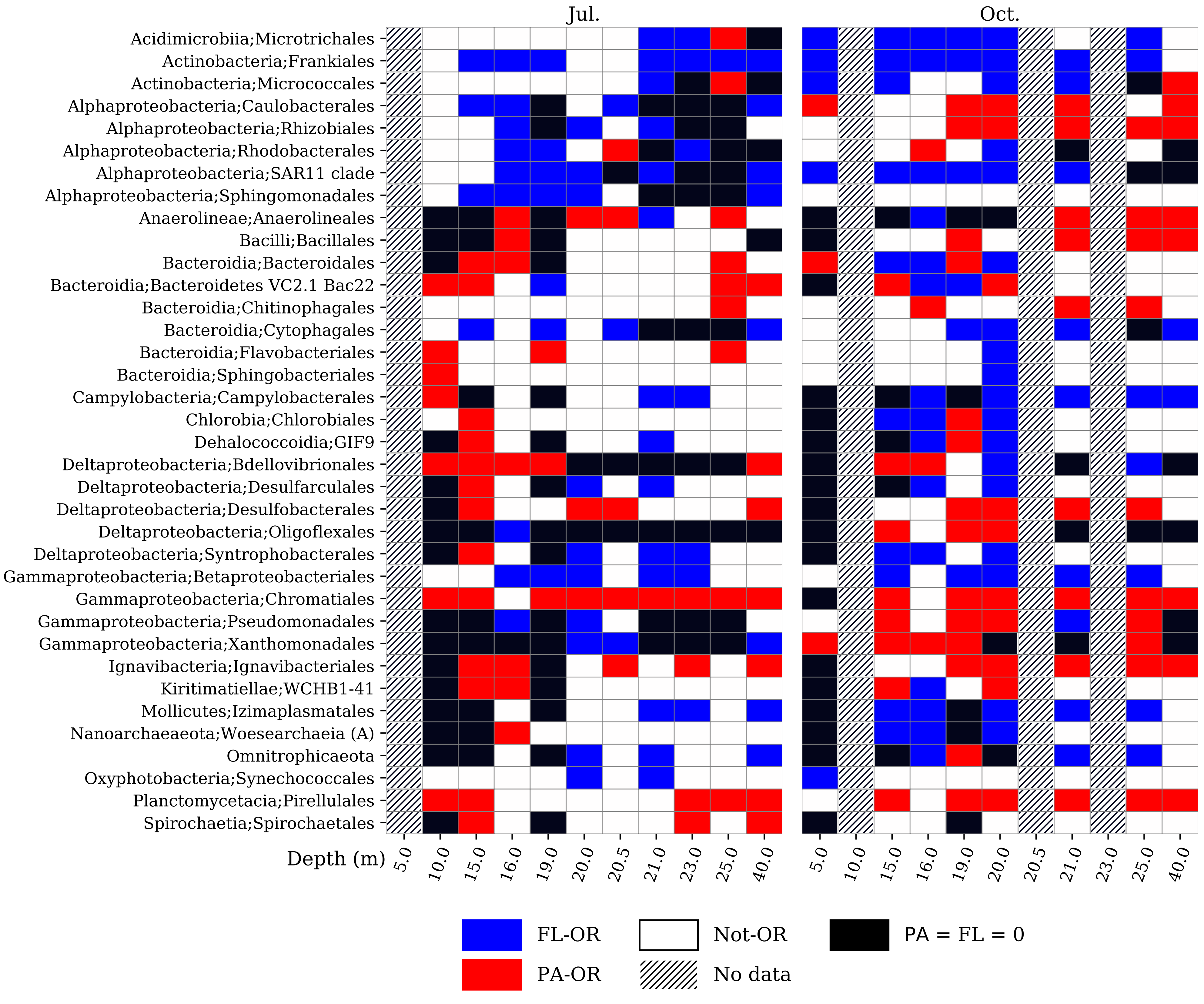

### CohenA_ORfactor_continuous_genuslevel.pdf

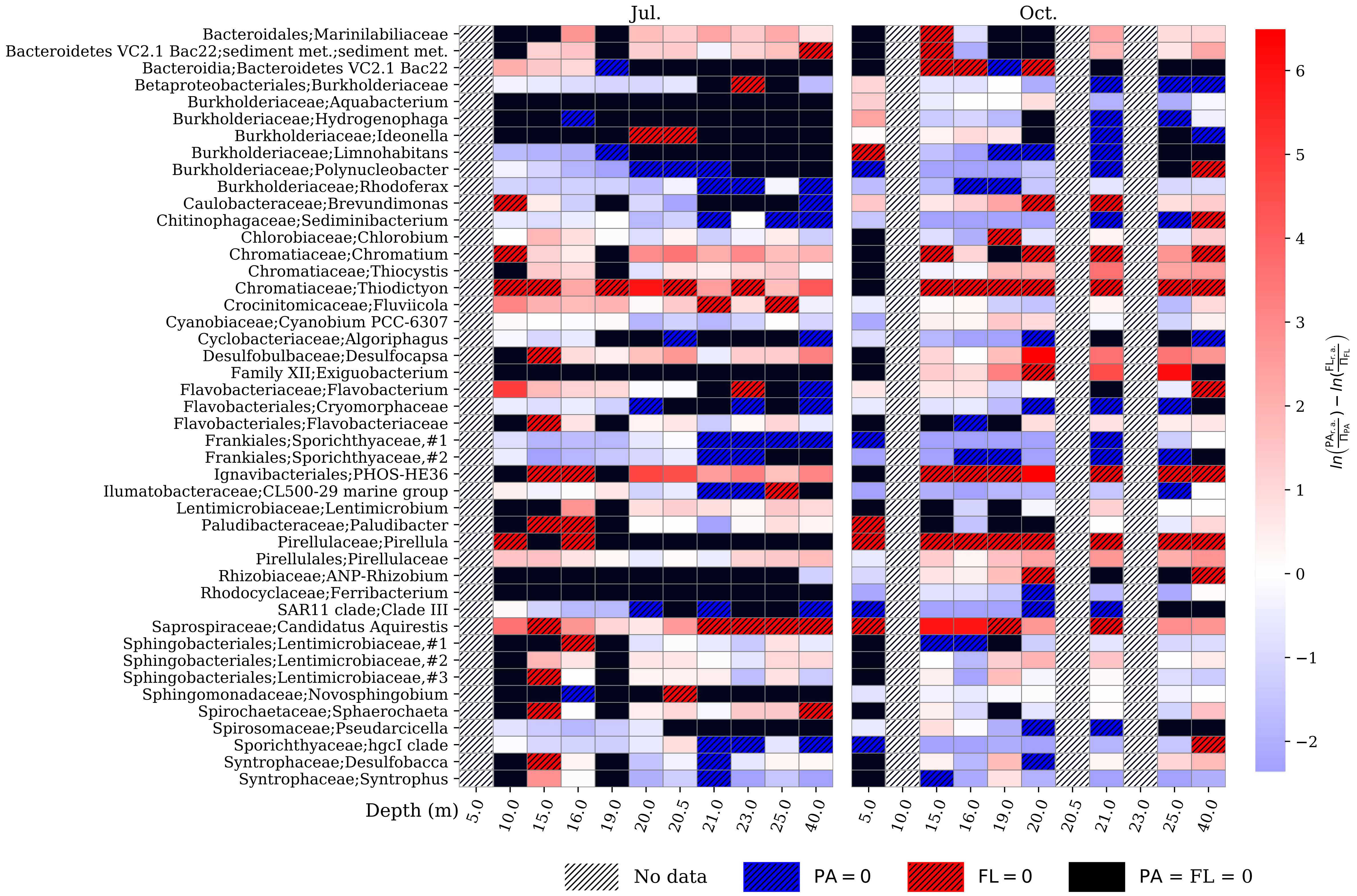

### CohenA_ORfactor_continuous_orderlevel.pdf

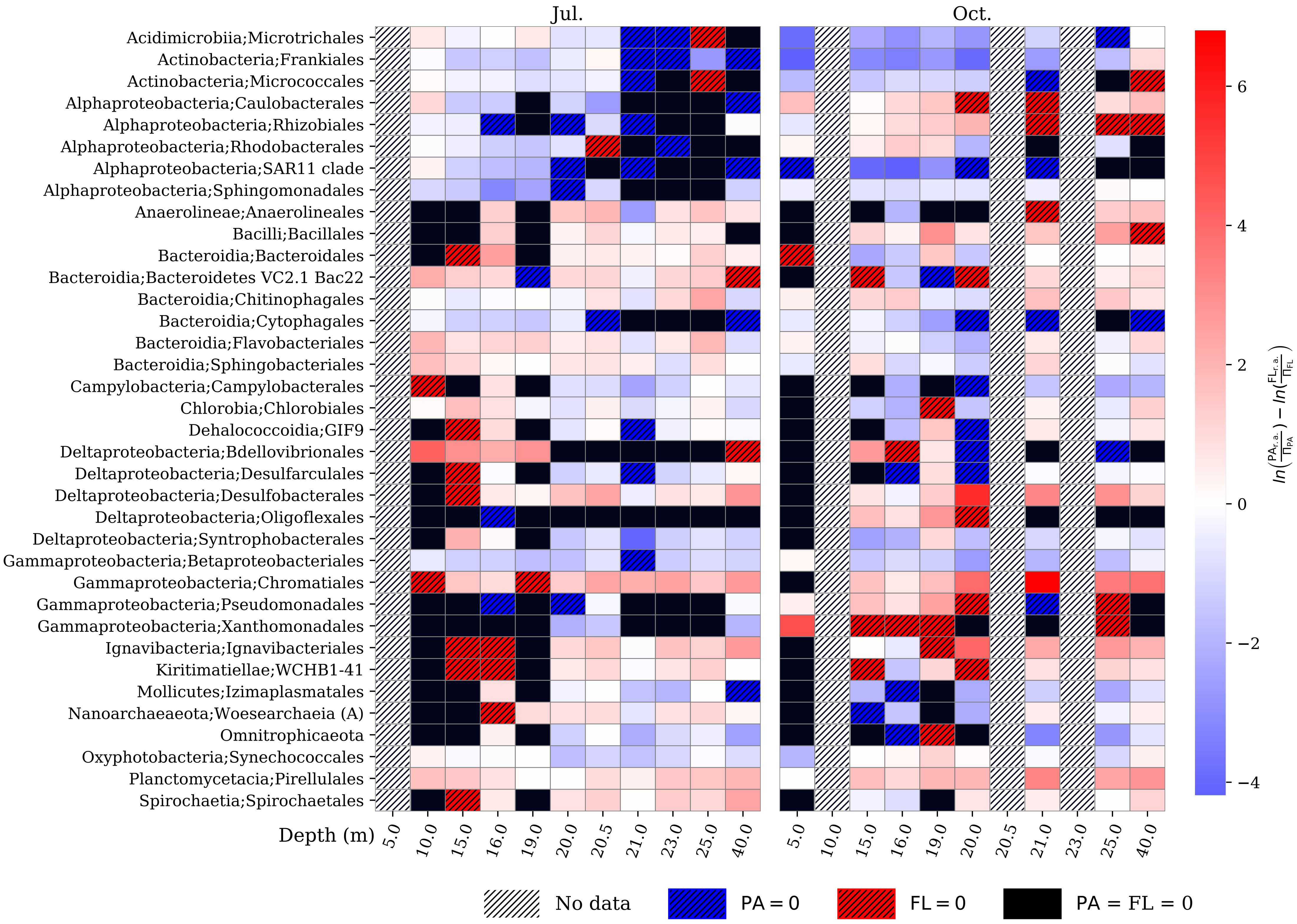

### CohenArelab_bubble_genuslevel.pdf

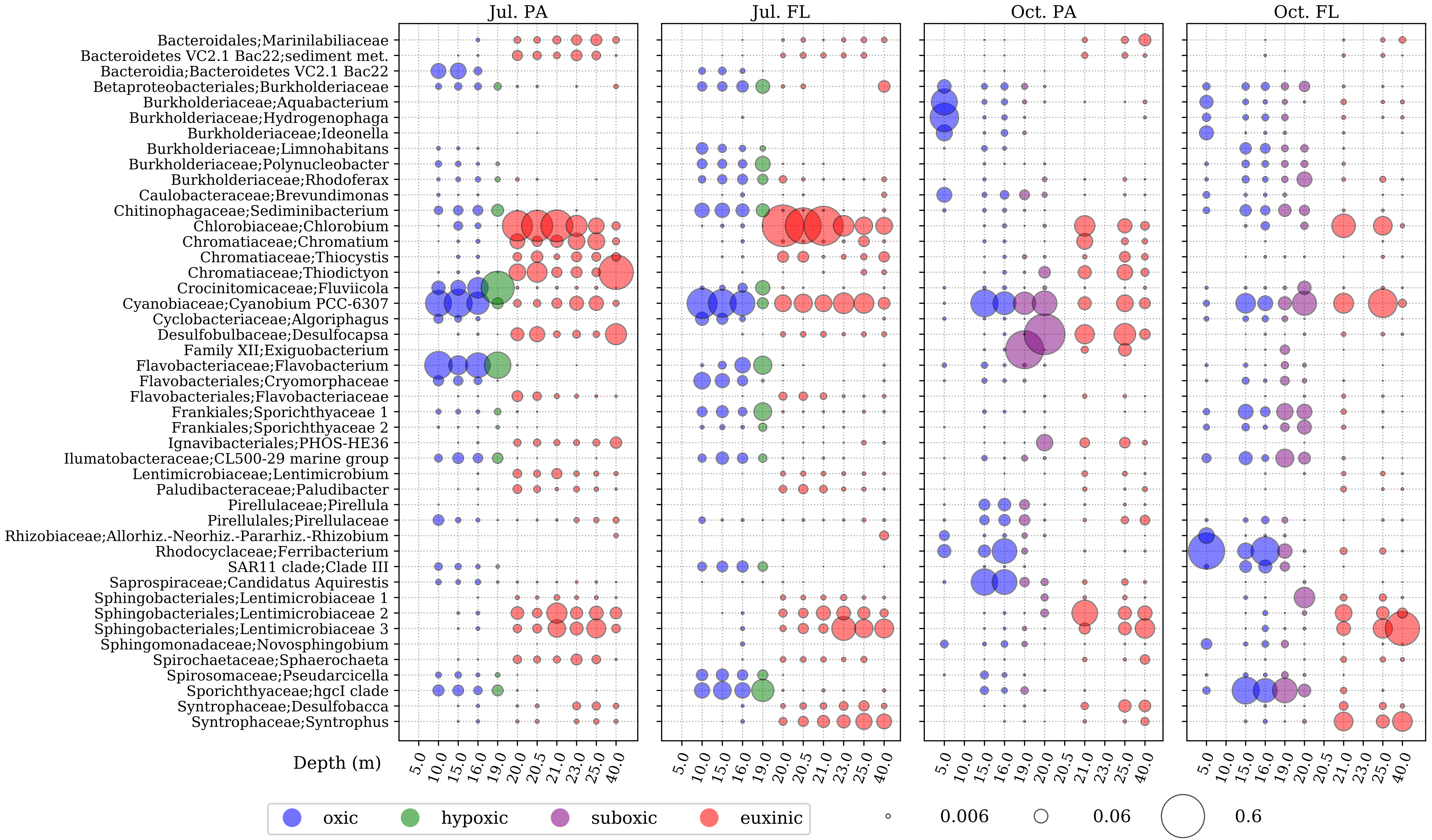

### CohenArelab_bubble_orderlevel.pdf

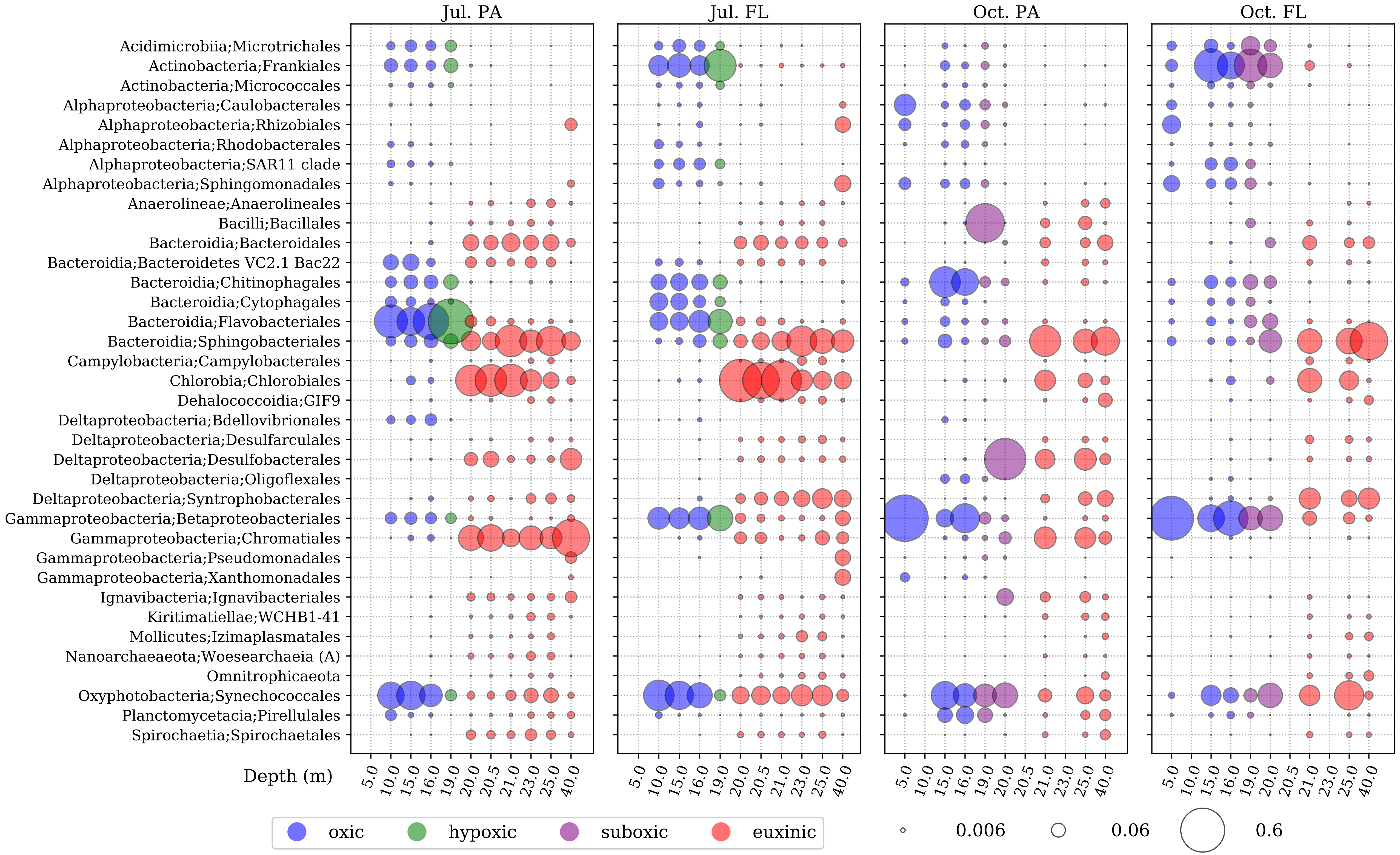
